# Predicting changes in bee assemblages following state transitions at North American dryland ecotones

**DOI:** 10.1101/746990

**Authors:** Melanie R. Kazenel, Karen W. Wright, Julieta Bettinelli, Terry L. Griswold, Kenneth D. Whitney, Jennifer A. Rudgers

**Author notes:** co-first authors (Melanie R. Kazenel, Karen W. Wright). corresponding author’s.

## Abstract

Drylands worldwide are experiencing ecosystem state transitions: the expansion of some ecosystem types at the expense of others. Bees in drylands are particularly abundant and diverse, with potential for large compositional differences and seasonal turnover across ecotones. To better understand how future ecosystem state transitions may influence bees, we compared bee assemblages and their seasonality among sites at the Sevilleta National Wildlife Refuge (NM, USA) that represent three dryland ecosystem types (and two ecotones) of the southwestern U.S. (Plains grassland, Chihuahuan Desert grassland, and Chihuahuan Desert shrubland). Using passive traps, we caught bees during two-week intervals from March – October, 2002 – 2014. The resulting dataset included 302 bee species and 56 genera. Bee abundance, composition, and diversity differed among ecosystems, indicating that future state transitions could alter bee assemblage composition in our system. We found strong seasonal bee species turnover, suggesting that bee phenological shifts may accompany state transitions. Common species drove the observed trends, and both specialist and generalist bee species were indicators of ecosystem types or months; these species could be sentinels of community-wide responses to future shifts. Our work suggests that predicting the consequences of global change for bee assemblages requires accounting for both within-year and among-ecosystem variation.

## Introduction

Drylands worldwide are experiencing ecosystem state transitions: the expansion of some ecosystem types at the expense of others^1,2^. These transitions include encroachment of C_3_ shrubland into C_4_ grassland^3^ and conversion of woodland to savanna^4^. It is through these transitions that the largest changes in dryland ecosystem processes are occurring^5–7^. State transitions can produce dramatic changes in carbon fluxes^8,9^, nutrient dynamics^10,11^, spatial heterogeneity in vegetation^12^, and consumer community composition^13,14^. Because drylands cover ~45% of land area on Earth^15^ and support over 2 billion people^16^, understanding how much dryland ecosystems currently differ in community composition can help to predict changes in future communities — and the ecosystem services they provide — under state transitions.

Bees may serve as important bio-indicators of state transitions and sentinels of altered ecosystem services^17,18^. In drylands, bees are important pollinators of both wild plants and agricultural crops^19,20^, and are particularly abundant and diverse. North America’s highest bee diversity occurs in the southwestern U.S. and northwest Mexico, and 75% of the continent’s bee species are found in the western U.S.^21,22^. Relative to mesic ecosystems, drylands can also host higher proportions of specialist bee species, which pollinate one or a few closely related plant species^23^. For example, creosote bush (*Larrea tridentata* (DC.) Coville), a widespread and abundant shrub in North American warm deserts^24^, is visited by 22 documented specialist bee species^25^. Cacti also host many specialists^26^. Communities dominated by specialist bees may be less resilient to state changes or pollinator declines than communities dominated by generalist bees, which can buffer plants against crashes in other bee species^27,28^. Future ecosystem state transitions could therefore substantially influence bees in drylands, making it important to understand potential vulnerabilities of dryland bee assemblages to these shifts.

Understanding variation in bee composition among habitat types can shed light on how ecosystem state transitions will influence bee assemblages. Prior studies have largely focused on bee assemblage variation within agricultural environments, along urban-rural gradients, or with habitat fragmentation^29,31^, while fewer studies have compared natural ecosystems. For instance, in Spain, shrub encroachment into grasslands corresponded with higher pollinator richness but fewer pollinator visits to forbs^32^. In xeric environments, some studies have documented bee species turnover across relatively small spatial scales^25,33,34^. For instance, during a single growing season, one study found lower bee abundance and richness in desert scrubland relative to riparian sites within a 4 km^2^ area in the Sonoran Desert^33^. In contrast, abundances of insect pollinator functional groups did not differ between creosote bush-dominated and adjacent annual forb-dominated microsites in the Mojave Desert^35^, although this study occurred on a smaller spatial scale with coarser taxonomic resolution. These contrasting results highlight the need for additional data to better understand the potential consequences for bee assemblages of specific state transitions in dryland ecosystems.

In addition, seasonal turnover in bee species composition suggests the potential for climate change to produce shifts in bee phenology^36^. Some bees may cue on climate variables for their emergence as adults, with temperature or precipitation conditions triggering the emergence of bee species at different times of year^37–39^. High temporal turnover in bee assemblage composition could thus indicate dominance of species with phenologies closely tied to climate, which may be particularly susceptible to phenological shifts under climate change. Understanding bee assemblage seasonality in ecosystem types predicted to expand or contract under climate change could thus be important for predicting bee assemblage responses to state transitions. However, while bee composition is well documented to vary seasonally within a community^40–42^, few studies have compared seasonal patterns among ecosystem types to discern how state transitions may shift bee phenology at the landscape scale within specific systems. Seasonal trends in bee abundance and richness were found to differ between natural and human-altered landscape types during a single year in California, USA^43^, and among agricultural land use classes during 3 years in New Hampshire, USA^44^. However, we lack studies that use long-term data to elucidate how general patterns of bee seasonality differ among natural ecosystem types that are expanding versus contracting.

This study compared bee assemblages and their seasonality among sites at the Sevilleta National Wildlife Refuge (NWR; NM, USA) representing three dryland ecosystem types of the southwestern U.S.: Chihuahuan Desert shrubland, Chihuahuan Desert grassland, and Plains grassland. Our sites occurred within a relatively small area (within 2–10 km of one another) that encompassed ecotones between the types, and shared the same regional pool of bee species. We used 13 years of monthly bee trap data to address two questions: (1) How much do bee assemblage abundance, composition, and diversity differ among sites representing major southwestern U.S. ecosystem types? (2) Do sites representing dryland ecosystem types differ in their degree of seasonal variation in bee abundance, composition, or diversity? We examined patterns among ecosystem types and months of the year by averaging across the time series, enabling us to identify general trends. Whereas this analysis focused on intra-annual and among-habitat variation in bee composition, a companion study will report inter-annual change over the time series, providing substantial additional complexity to the analysis. Forthcoming work will also examine the potential abiotic and biotic drivers of bee assemblage trends.

## Methods

### Ecosystem types

The Sevilleta NWR is located on the northern edge of the Chihuahuan Desert in central New Mexico, USA, and includes five ecosystem types that together represent ~80 million ha of the southwestern U.S. Total annual precipitation is ~250 mm, with ~60% occurring during the summer monsoon season from July through early September^45^. We focused on three major ecosystem types: Chihuahuan Desert shrubland, which is dominated by creosote bush (*Larrea tridentata*), Chihuahuan Desert grassland, which is dominated by black grama grass (*Bouteloua eriopoda* (Torr.) Torr.), and Plains grassland, which is dominated by blue grama grass (*Bouteloua gracilis* (Willd. Ex Kunth) Lag. Ex Griffiths) (Table 1). Transitions among these ecosystem types are predicted to occur under climate change, with Chihuahuan Desert shrubland encroaching upon Chihuahuan Desert grassland, which is predicted to replace Plains grassland^46–49^. In our study, the two Chihuahuan Desert sites were separated by ~2 km; the Plains grassland site was ~10 km from the Chihuahuan Desert sites (Table 1; Supplementary Fig. S1).

**Table 1.** Latitude, longitude, and elevation of study sites representing three ecosystem types of the southwestern U.S., along with current versus predicted future foundation species.

| Ecosystem type | Latitude | Longitude | Elevation<br>(m) | Current<br>foundation<br>species | Future<br>foundation<br>species |
| --- | --- | --- | --- | --- | --- |
| Desert shrubland | 34.3329 | -106.7358 | 1615 | Creosote bush | Creosote bush |
| Desert grassland | 34.3362 | -106.7212 | 1616 | Black grama | Creosote bush |
| Plains grassland | 34.3364 | -106.6345 | 1670 | Blue grama | Black grama |

### Bee collection

Bees were sampled along five transects located within each of the three focal ecosystem types (Supplementary Fig. S1). To sample bees, we installed one passive funnel trap at each end of five 200 m transects/site. As bees are capable of movement between traps within a site, our traps represent non-independent sampling locations, which we accounted for in our statistical analyses (see below). Each trap consisted of a 946 mL paint can filled with ~275 mL of propylene glycol and topped with a plastic automotive funnel with the narrow part of the funnel sawed off (funnel height = 10 cm, top diameter = 14 cm, bottom diameter = 2.5 cm; Supplementary Fig. S2). The funnels’ interiors were painted with either blue or yellow fluorescent paint (Krylon, Cleveland, OH or Ace Hardware, Oak Brook, IL). On each transect, we randomly assigned one trap to be blue and the other to be yellow (total across the three sites: *N* = 30 traps, with 15 traps/color). Because different bee taxa are known to be attracted to blue versus yellow^50^, we summed the samples collected in the two traps on a given transect. Each trap was placed on a 45 cm high platform that was surrounded by a 60 cm high chicken wire cage to prevent wildlife and wind disturbance (Supplementary Fig. S2). Funnel traps provide a measure of bee activity, not a measure of presence, and may be biased by bee taxon, sociality, sex, pollen specialization, floral resource availability, and microsite conditions^50–53^. From 2002 to 2014, bees were sampled each month from March through October. Traps were opened each March as close as possible to the first day of spring, and left open for 14 d, after which the bee specimens were collected. The traps were then closed for 14 d. This two-week cycle was repeated through October. Bees were rinsed and stored in 70% ethanol until processed.

### Bee identification

Bees were identified to species by K.W.W. and T.L.G. Certain groups of bees could not be identified to species, either because there are no practicing experts in the bee group and species are unnamed for our study region, or because there are no revisions within the bee group to separate named from unnamed species. In these cases, we separated females into morphotypes as best as possible. The males of these groups could not be reliably linked to the females and were therefore excluded from the dataset. The major groups treated in this manner were the genera *Sphecodes, Protandrena*, and *Nomada*, the subgenera *Dialictus* and *Evylaeus* of the genus *Lasioglossum*, and the subgenus *Micrandrena* of *Andrena*. We excluded *Nomada* from our analyses due to low abundance and lack of ability to distinguish among species. New species of relatively well-known genera were recognized, and the qualifier aff. was used with uncertain identifications. Voucher specimens were deposited at the University of New Mexico’s Museum of Southwestern Biology and the USDA-ARS Pollinating Insects Research Unit’s U.S. National Pollinating Insects Collection. Information related to these specimens is available via the Symbiota Collections of Arthropods Network (https://scan-bugs.org).

### Analysis

#### Dataset

We created a species matrix in which cells contained the mean abundance of each bee species for each month of collection, averaged over the years of collection (2002 – 2014). Each row was a unique trapping transect, with five transects per ecosystem type per month (*N* = 120 observations). Means were calculated using the <reshape2> package^54^ in R version 3.4.2^55^. To examine whether assemblage-level patterns were driven by common or rare species, we ran all abundance, composition, and diversity analyses (described below) on the full dataset, on a dataset with singleton bee species (those caught only on a single transect, in a single month) removed, and finally on a subset of the dataset containing only the bee species that were present in >5% of the samples.

#### Overview

Analyses addressed our two key questions within one set of statistical models (described below). First, (1) How much do bee assemblage abundance, composition, and diversity differ among sites representing major southwestern U.S. ecosystem types? was determined by the statistical significance and magnitude of the effect of ecosystem type in our models. We also compared the effect size of ecosystem type against the effect size of month of sampling to estimate the relative importance of inter-ecosystem versus seasonal variability. Then, to address (2) Do sites representing dryland ecosystem types differ in their degree of seasonal variation in bee abundance, composition, or diversity? we evaluated whether the interaction between ecosystem type and month of sampling was statistically significant, indicating that ecosystems differed in the seasonality of bee abundance, composition, or diversity.

#### Bee assemblage composition and turnover

For bee composition, we calculated Bray-Curtis similarities in Primer version 6.1.13^56^. We then tested for the influence of ecosystem type, month of sampling, and the random effect of transect, which was nested within ecosystem type to account for the repeated measures design, using perMANOVA (version 1.0.3) with 9999 permutations of residuals under a reduced model. We additionally examined whether ecosystem types or months differed in bee assemblage dispersion using permDISP in Primer^56^. We visualized assemblage composition with non-metric multidimensional scaling analysis (NMDS) implemented with 500 restarts in Primer. For each ecosystem type, we assessed bee species turnover among months, as well as the rate of community change, using the <codyn> package in R^57^. Finally, to identify which taxa contributed most to bee assemblage (i) divergence among ecosystem types and (ii) divergence among months within each ecosystem type, we calculated Dufrene-Legendre (DL) indicator species values using the indval function in the <labdsv> R package^58^, which takes both species’ presence/absence and abundance into account.

#### Bee diversity and abundance

For bee diversity, we calculated the Shannon diversity index (*H’*), species richness, and evenness (Pielou’s *J*) using the <vegan> package in R^59^. We then used linear mixed effects models to examine the influences of ecosystem type, sampling month, and their interaction (fixed effects), as well as transect identity (random effect nested within ecosystem type), on these three responses, as well as on total bee abundance (function lmer, <lme4> package in R)^60^. When there was a significant ecosystem type x sampling month interaction, we tested *a priori* contrasts for pairs of months within each ecosystem type and for pairs of ecosystem types within each month using Tukey-Kramer multiple comparisons in the <emmeans> package in R^61^.

## Results

### The dataset

We captured a total of 70,951 individuals representing 302 species during the 13 years of monthly trapping (see Supplementary Table S1 for a full species list). Species were distributed across 6 families and 56 genera (Supplementary Table S1 and Fig. S3). Our dataset was dominated by a small number of abundant species and contained a large number of rare species (Supplementary Fig. S4). The most commonly collected species were *Lasioglossum semicaeruleum* (36% of all collected specimens), *Agapostemon angelicus* (21%), *Diadasia rinconis* (7%), *Melissodes tristis* (5%), *Anthophora affabilis* (5%), and *Eucera lycii* (3%). Amongst the collected species, 30% were singletons, and 58% were found in <5% of all samples.

### Bee assemblage composition: temporal variation surpassed differences among sites representing dryland ecosystem types

#### Variation among ecosystems

All ecosystems significantly diverged in bee assemblage composition, and this pattern was present during all months (Table 2, Fig. 1). The greatest difference among ecosystems occurred in October, when the Plains grassland bee assemblage diverged most strongly from the Chihuahuan Desert shrubland (mean similarity: 41.4, *P* = 0.0089) and also diverged from the Chihuahuan Desert grassland (mean similarity: 51.6, *P* = 0.0080). The three ecosystem types did not differ in assemblage dispersion (*F*_2,117_ = 0.52, *P* = 0.71), indicating similar levels of temporal beta-diversity among ecosystem types (Fig. 1).

**Figure 1.**
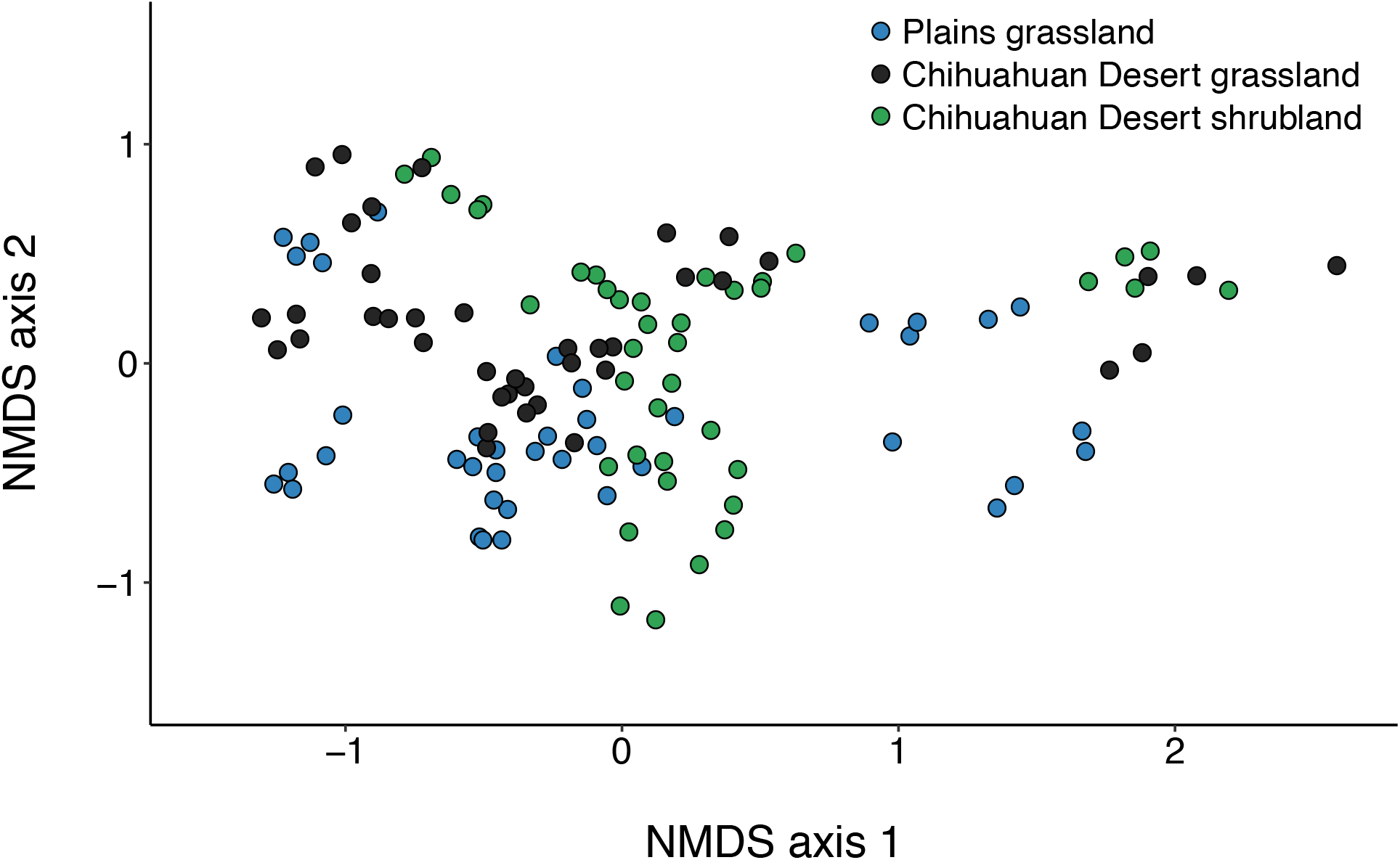
Non-metric multidimensional scaling (NMDS) plot depicting variation in bee species composition among sites representing three dryland ecosystem types: Plains grassland (blue points), Chihuahuan Desert grassland (black points), and Chihuahuan Desert shrubland (green points). NMDS was run with 500 randomized re-starts and 2D stress = 0.13. On average, all ecosystem types significantly differed from one another (Table 2): Plains grassland versus Chihuahuan Desert grassland (*P* = 0.0082), Plains grassland versus Chihuahuan Desert shrubland (*P* = 0.0075), and Chihuahuan Desert grassland versus shrubland (*P* = 0.0084).

**Table 2.** Results of 1) perMANOVA with 9999 permutations to test for the influence of ecosystem type and month of sample collection on bee assemblage composition, using a Bray-Curtis similarity metric, and 2) permDISP examining differences among ecosystem types and months in bee assemblage dispersion.

|  | *num. df* | perMANOVA |  |  |  | permDISP |  |  |
| --- | --- | --- | --- | --- | --- | --- | --- | --- |
|  |  | *SS* | *MS* | *pseudo-F* | *P* | *denom. df* | *F* | *P* |
| Ecosystem | 2 | 23252.00 | 11626.00 | 34.54 | 0.0001 | 117 | 0.52 | 0.7074 |
| Month | 7 | 139860.00 | 19981.00 | 94.02 | 0.0001 | 112 | 5.92 | 0.0002 |
| Ecosystem x month | 14 | 34139.00 | 2438.50 | 11.48 | 0.0001 |  |  |  |
| Transect (ecosystem) | 12 | 4039.70 | 336.64 | 1.58 | 0.0003 |  |  |  |
| Residuals | 84 | 17851.00 | 212.51 |  |  |  |  |  |

#### Indicators of variation among ecosystems

We identified 43 bee species as ecosystem indicators according to their Dufrene-Legendre (DL) indicator species values (Table 3). Of these, 21 species were indicators of Chihuahuan Desert shrubland, 14 species were indicators of Plains grassland, and 8 species were indicators of Chihuahuan Desert grassland. All three ecosystems had indicator species within the families Andrenidae, Apidae, Halictidae, and Megachilidae, and one Plains grassland indicator species was in the family Colletidae (Table 3, Fig. 2). In all three ecosystems, *Lasioglossum semicaeruleum* (an indicator of the Desert grassland), *Agapostemon angelicus* (an indicator of Plains grassland)*, Diadasia rinconis*, and *Melissodes tristis* were among the five most abundant bee species (Fig. 2). *Anthophora affabilis* was also within the five most abundant species in the Plains and Chihuahuan Desert grasslands, while *Perdita larreae* (a creosote bush specialist) was abundant in and an indicator of the Chihuahuan Desert shrubland (Fig. 2).

**Figure 2.**
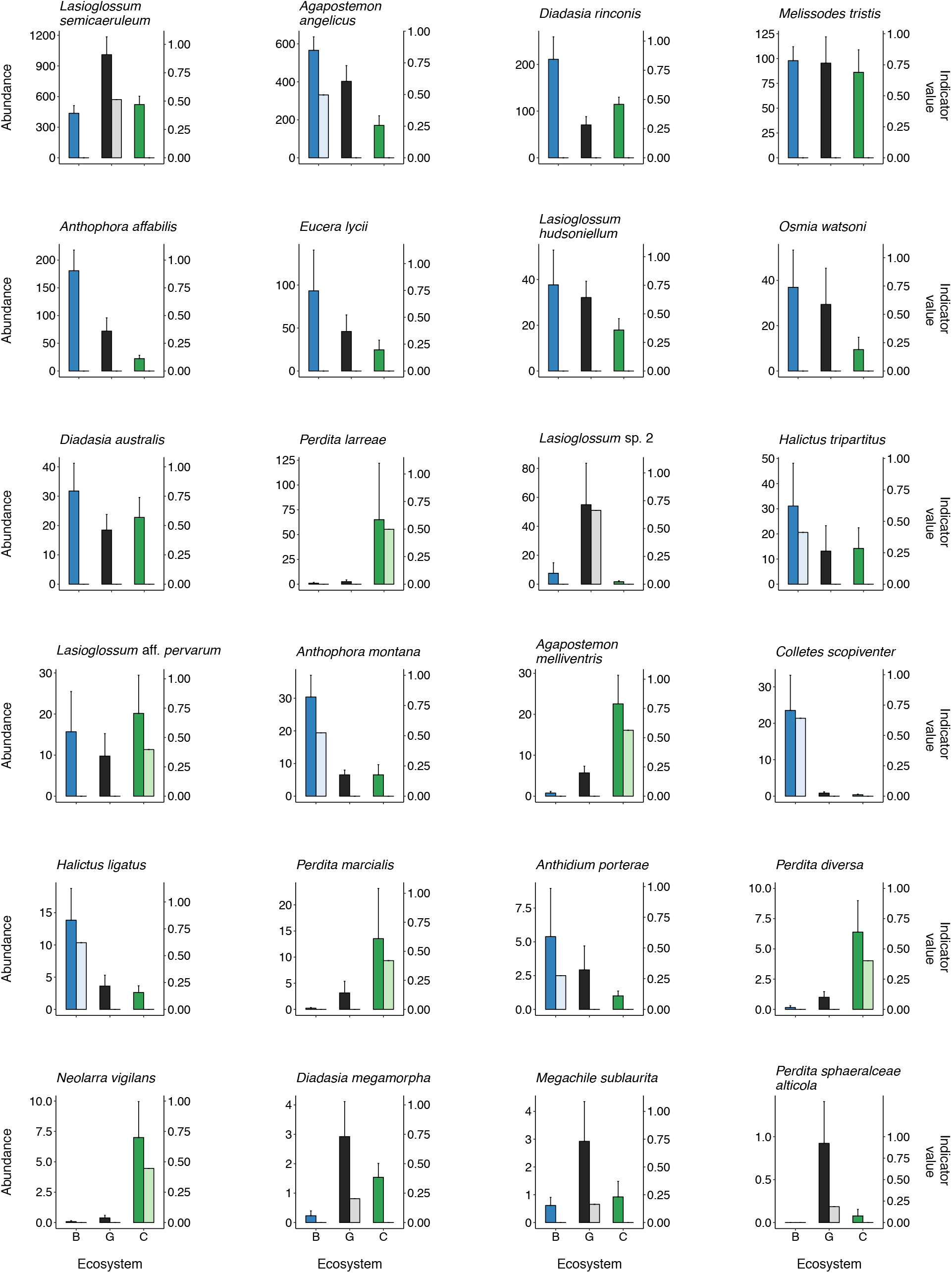
Mean yearly abundance + s.e. (darker, leftmost bar in each pair) and Dufrene-Legendre (DL) indicator species value (lighter, rightmost bar in each pair) for important bee species within each ecosystem type (Plains grassland, blue bars, B; Chihuahuan Desert grassland, black bars, G; Chihuahuan Desert shrubland, green bars, C). Included bee species were within the 20 most abundant species found across the study, and/or were top indicator species of particular ecosystem types according to DL indicator value. Plots are arranged from left to right by mean yearly abundance across ecosystem types.

**Table 3.**
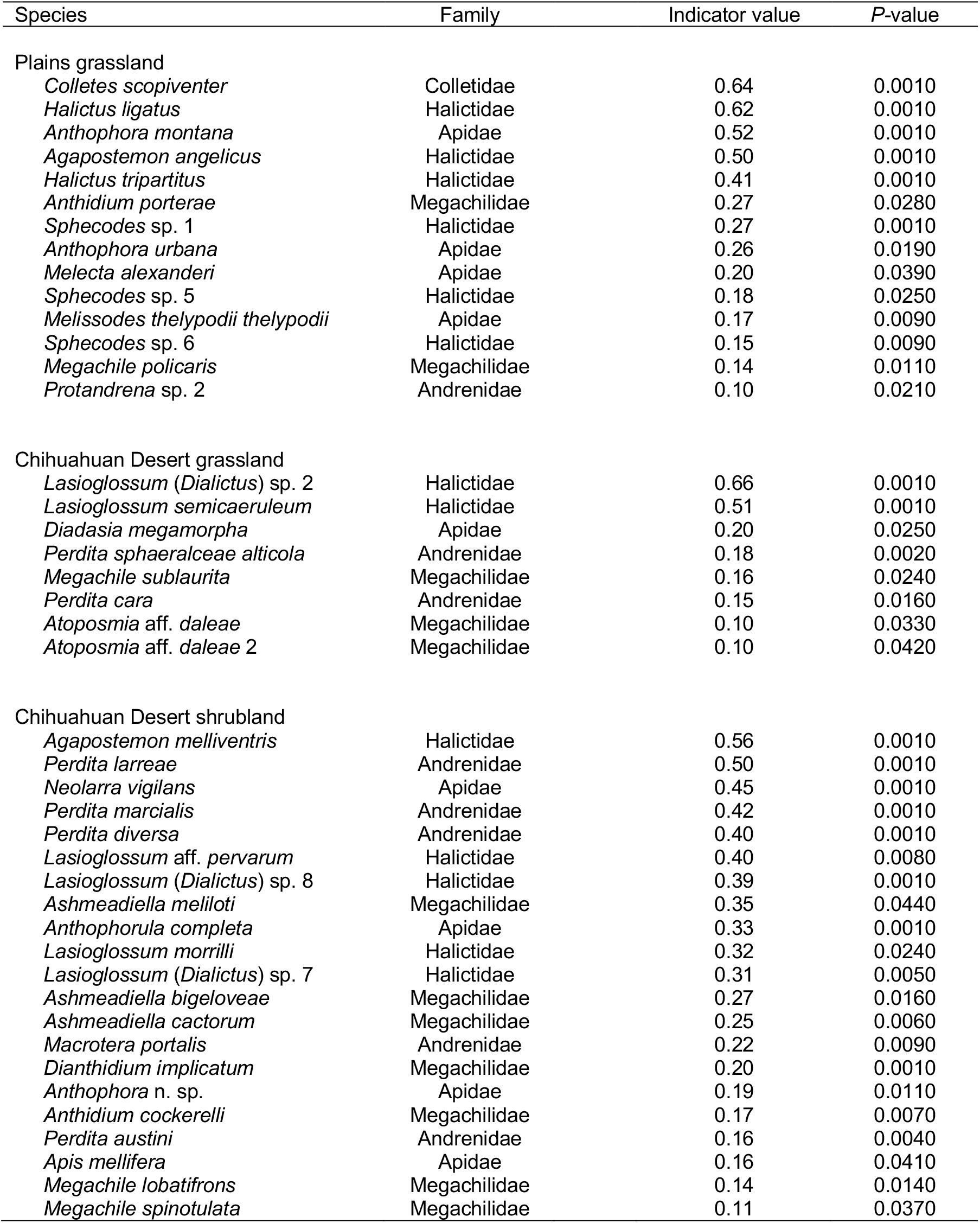
Indicator species for each ecosystem (Plains grassland, Chihuahuan Desert grassland, and Chihuahuan Desert shrubland) according to Dufrene-Legendre indicator species value. Species are listed from highest to lowest indicator value within each ecosystem.

#### Temporal variation

The month of sample collection explained an order of magnitude more variation in bee assemblage composition than did ecosystem type (Table 2, Fig. 3). Generally, assemblages diverged between the early and late months of the year and converged during the middle of the summer. Across ecosystems, the pair of months most divergent in bee composition was March versus October (mean similarity = 12.7, *P* = 0.0001). In contrast, June and July were most similar in bee composition (mean similarity = 64.0, *P* = 0.0001).

**Figure 3.**
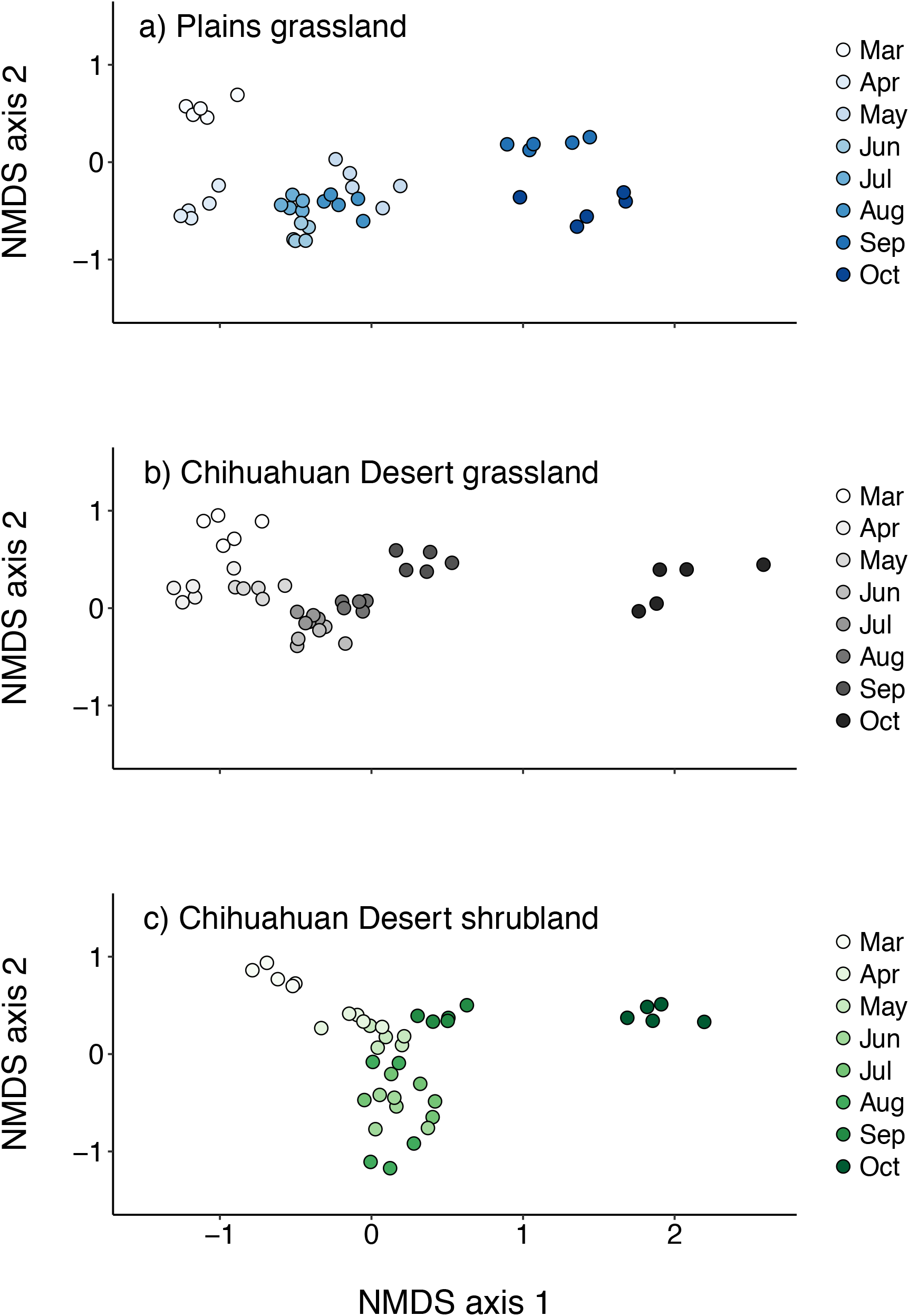
Non-metric multidimensional scaling (NMDS) plots depicting variation in bee species composition among months for sites representing three dryland ecosystems: (a) Plains grassland, (b) Chihuahuan Desert grassland, and (c) Chihuahuan Desert shrubland. NMDS was run with all samples together, with 500 randomized re-starts and 2D stress = 0.13.

Months additionally differed from one another in the magnitude of assemblage dispersion, a metric that captures the degree of beta-diversity across both sites and transects (Table 2, Fig. 3). The strongest differences in beta-diversity were between March or June, which had the smallest multivariate dispersions (mean ± s.e., March: 21.0 ± 1.5, June: 20.4 ± 0.8), against October, which had the largest average dispersion across ecosystems (29.8 ± 1.8).

### Bee abundance and diversity: temporal variation exceeded variation among sites representing dryland ecosystem types

#### Abundance

As with composition, across months, ecosystems diverged significantly from one another in total bee abundance (Table 4). Bee abundance was on average 43% lower in the Chihuahuan Desert shrubland relative to the Desert and Plains grassland sites, respectively, from March through July (Fig. 4a). However, abundances within the ecosystems converged in August, and abundance differences disappeared in September and October (Fig. 4a), as indicated by a significant interaction between ecosystem type and month of collection (Table 4: Ecosystem x Month, *P* < 0.0001).

**Figure 4.**
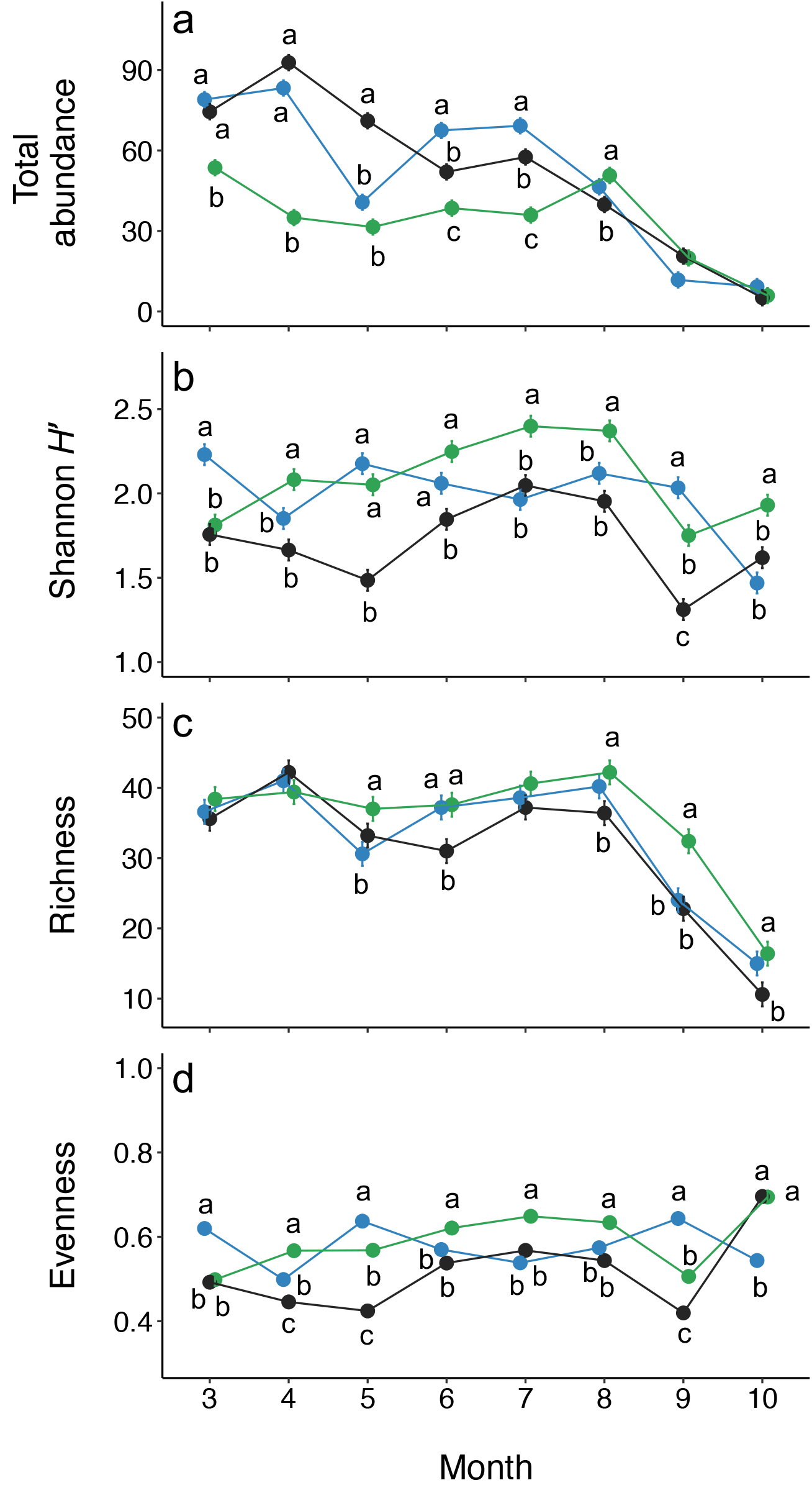
Variation across sampling months in per-transect bee abundance and diversity (± s.e.) as measured by a) total bee abundance, b) Shannon diversity index (*H’*), c) richness, and d) evenness (Pielou’s *J*) for sites representing three dryland ecosystem types: Plains grassland (blue points), Chihuahuan Desert grassland (black points), and Chihuahuan Desert shrubland (green points). Letters denote contrasts between ecosystems within a given month; ecosystems labeled with different letters differed significantly from one another in the relevant abundance/diversity metric. Points lacking letters did not differ significantly from any other ecosystem in the given month. For total abundance, s.e. values were <0.1.

**Table 4.** Results of linear mixed effects models testing the influences of ecosystem type and month of sample collection on total bee abundance, as well as bee assemblage Shannon diversity index (*H*’), richness, and evenness (Pielou’s *J*).

| | *df* | Total abundance | | Shannon $H'$ | | Richness | | Evenness | |
| --- | --- | --- | --- | --- | --- | --- | --- | --- | --- |
| | | $\chi^2$ | *P* | $\chi^2$ | *P* | $\chi^2$ | *P* | $\chi^2$ | *P* |
| Ecosystem | 2 | 45.19 | < 0.0001 | 34.72 | < 0.0001 | 1.38 | 0.50044 | 47.28 | < 0.0001 |
| Month | 7 | 796.09 | < 0.0001 | 105.64 | < 0.0001 | 221.14 | < 0.0001 | 85.33 | < 0.0001 |
| Ecosystem x month | 14 | 359.31 | < 0.0001 | 142.78 | < 0.0001 | 28.13 | 0.01368 | 320.07 | < 0.0001 |

#### Diversity

Ecosystems also diverged in bee diversity as measured by the Shannon index and Pielou’s evenness (Table 4). Differences in Shannon diversity (Fig. 4b) among ecosystems were more strongly driven by evenness (Fig. 4d) than by richness (Fig. 4c). On average across all months of sampling, the Chihuahuan Desert shrubland ecosystem had the highest bee Shannon diversity and evenness, with these diversity metrics 5% (Shannon diversity) and 2% (evenness) higher than in the Plains grassland. In turn, Plains grassland diversity metrics were 16% (Shannon diversity) and 12% (evenness) higher than the Chihuahuan Desert grassland. In contrast, on average across months, the ecosystems did not significantly differ in bee species richness (Table 4, Fig. 4c).

Importantly, differences among ecosystems in all diversity metrics varied by month of the year (Fig. 4, Table 4: Ecosystem x Month – Shannon diversity: *P* < 0.0001, richness: *P* = 0.0137, evenness: *P* < 0.0001), indicating that dryland ecosystem types differed in their degree of seasonal variation in bee diversity (Question 2). Specifically, Shannon diversity was greater in the Chihuahuan Desert shrubland than in the Desert grassland in all months except for March; differences in Shannon diversity were largest in May and September, when Shannon diversity respectively was 38% and 33% higher in the Chihuahuan Desert shrubland relative to grassland (Fig. 4b). Shannon diversity was also higher in the Chihuahuan Desert shrubland relative to Plains grassland in April, July, August, and October (Fig. 4b). The largest difference occurred in October, in which Shannon diversity was 31% higher in the Chihuahuan Desert shrubland than Plains grassland. However, this trend was reversed in both March and September, when Shannon diversity was 19% and 16% higher, respectively, in Plains grassland than in shrubland (Fig. 4b). The two grassland ecosystems differed in Shannon diversity in March, May, June, and September, with greater Shannon diversity in the Plains relative to Chihuahuan Desert grassland in all of these months (Fig. 4b).

### Sites representing dryland ecosystems diverged in the magnitude of seasonal variation in bee assemblage composition, abundance, and diversity

#### Assemblage composition

Bee assemblage composition varied strongly among months, with the magnitude of seasonal change differing among ecosystems (Figs. 3,5; Table 2: Ecosystem X Month, *P* = 0.0001). The Chihuahuan Desert grassland had the greatest seasonal turnover in bee species composition (Fig. 3b), and the highest rate of compositional change from month to month (Fig. 6). In contrast, the Chihuahuan Desert shrubland had the lowest seasonal composition change (Figs. 3c,6), with low turnover between July and August, and between August and September, compared to the other ecosystems (Fig. 5). Among months, in all ecosystem types, bee species composition differed most strongly between March and either September (Plains grassland: mean similarity = 16.4, *P* = 0.0077) or October (Desert grassland: mean similarity = 9.2, *P* = 0.0091; shrubland: mean similarity = 11.7, *P* = 0.0070) (Fig. 3). In contrast, in all ecosystems, June and July were most compositionally similar to one another, with low turnover between them (Figs. 3,5; Plains grassland: mean similarity = 76.2, *P* = 0.0091; Desert grassland: mean = 72.42, *P* = 0.0077; Desert shrubland mean = 70.4, *P* = 0.0156). Seasonal patterns in bee assemblage composition were largely driven by common rather than rare species, as indicated by very few qualitative differences in analysis outcomes when excluding singletons or moderately rare species (see Supplementary Fig. S5).

**Figure 5.**
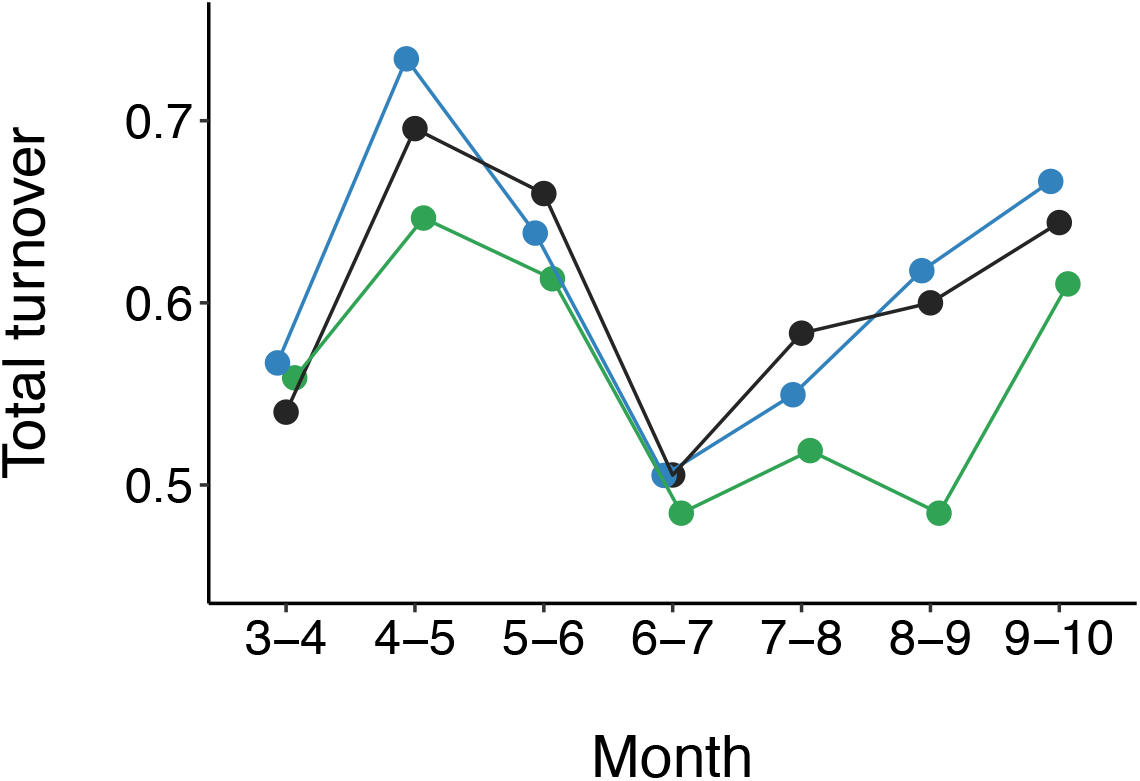
Total bee species turnover between pairs of months (indicated on the x-axis) for sites representing three dryland ecosystem types: Plains grassland (blue points), Chihuahuan Desert grassland (black points), and Chihuahuan Desert shrubland (green points).

**Figure 6.**
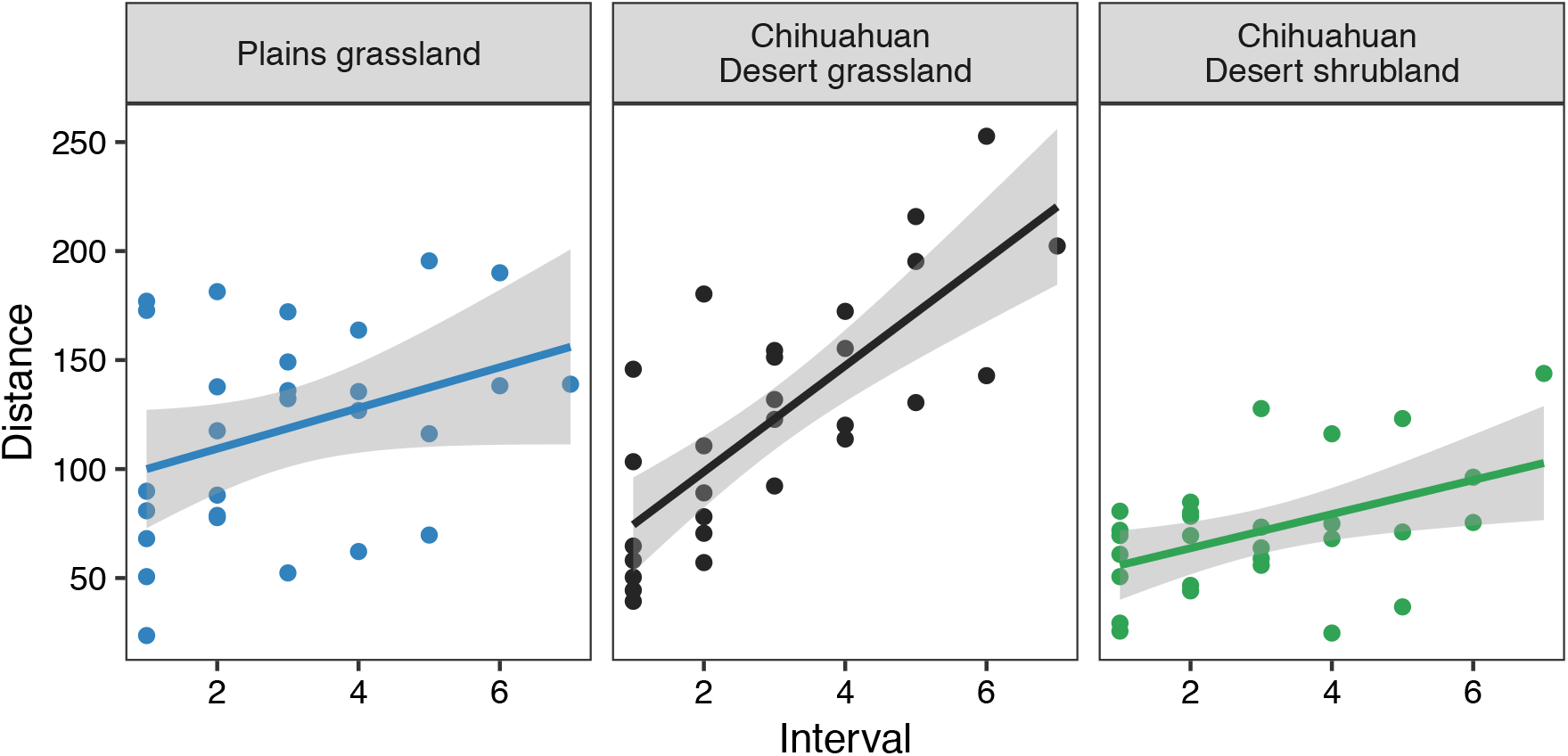
Average rate of bee assemblage change during March through October in sites representing three ecosystem types: Plains grassland (slope = 9.34, s.e. = 4.99, *t* = 1.9, *P* = 0.0725), Chihuahuan Desert grassland (slope = 24.32, s.e. = 3.99, *t* = 6.1, *P* < 0.0001), and Chihuahuan Desert shrubland (slope = 7.81, s.e. = 2.92, *t* = 2.7, *P* = 0.0128). Intervals (x-axis) represent time lags between all pairwise combinations of months. Distances (y-axis) correspond with differences in bee assemblage composition between pairs of months, calculated as Euclidean distances. The slope of each line indicates the rate of bee assemblage change in each ecosystem.

#### Abundance

Like species composition, total bee abundance also varied seasonally across the three ecosystem types (Table 4), and ecosystem types exhibited differing trends in total abundance over the course of the season (Fig. 4a). In the Chihuahuan Desert grassland, bee abundance increased from March to April, then generally declined through the rest of the season (Fig. 4a). In contrast, the Plains grassland had similar levels of bee abundance in March and April (*df* = 84, *t* = −1.13, *P* = 0.95), followed by a ~50% decrease in abundance between April and May (*df* = 84, *t* = 11.12, *P* < 0.0001) and a 66% increase in abundance between May and June (*df* = 84, *t* = −6.99, *P* < 0.0001). Between July and August, while bee abundance decreased ~30% within both the Chihuahuan Desert grassland (*df* = 84, *t* = 4.62, *P* = 0.0004) and Plains grassland ecosystems (*df* = 84, *t* = 5.93, *P* < 0.0001), it increased by 40% within the Chihuahuan Desert shrubland (*df* = 84, *t* = −3.86, *P* = 0.0053) (Fig. 4a). Across ecosystem types, bee abundances were generally lower in September and October relative to all other months (Fig. 4a).

#### Diversity

Within each ecosystem, most months had similar levels of species richness, with some exceptions (Fig. 4c). Notably, there was a sharp decline in richness between August and October across all three ecosystems (Fig. 4c). During this period, richness declined by 70% within the Chihuahuan Desert grassland (*df* = 84, *t* = 11.18, *P* < 0.0001) and by 60% within both the Plains grassland (*df* = 84, *t* = 10.92, *P* < 0.0001) and Desert shrubland (*df* = 84, *t* = 11.18, *P* < 0.0001). However, month-to-month trends in Shannon diversity and evenness diverged among ecosystems (Figs. 4b,d). Patterns in total abundance, Shannon diversity, richness, and evenness were all largely driven by common rather than rare species (see Supplementary Fig. S6).

#### Indicators of temporal variation within ecosystem types

Certain bee taxa were indicators of specific months across all three ecosystems according to their DL indicator values (Supplementary Tables S3-S6). These included *Osmia* species, *Eucera lycii, Anthophora porterae*, and *Melecta pacifica* (March), *Dioxys* and *Anthophora* species (April), *Diadasia australis* and *rinconis* (June), *Martinapis lutericornis, Halictus ligatus*, and *Melissodes tristis* (July), and *Perdita semicaerulea* and *marcialis* (August) (Supplementary Tables S3-S6). September and October lacked indicator species shared by all ecosystems.

In contrast, certain bee taxa were only characteristic of a given month within one or two ecosystems (Supplementary Tables S3-S6). For instance, in June, the shrubland site had 5 indicator species in the genus *Lasioglossum*; one of these was also characteristic of the desert grassland site (Supplementary Tables S3,S5,S6). In July, *Perdita* species (Andrenidae) were indicators of the Chihuahuan Desert sites but not the Plains site, while the Chihuahuan Desert and Plains grassland sites had indicator species in the Halictidae (especially *Lasioglossum*), and differing members of the Apidae were characteristic of different ecosystems (Supplementary Tables S3-S6). In September, *Perdita* species (Andrenidae) were characteristic of the Plains grassland, *Macrotera* (Andrenidae) were characteristic of the desert sites, and differing members of the Apidae were characteristic of each site (Supplementary Tables S3-S6). Finally, 25 species were indicators of a particular month in one or two ecosystem types, and were then indicators of a different month, often the following one, in the other ecosystem(s) (Supplementary Table S3).

## Discussion

### Ecosystem state transitions: potential consequences for bee assemblages

We found large variation in bee assemblages and their seasonality among sites representing three dryland ecosystem types of the southwestern U.S. These results indicate the potential for future ecosystem state transitions to alter bee assemblage composition in our dryland system. Overall, ecosystem types in our study had similar levels of bee species richness but differed from one another in species evenness and composition. These results imply that state transitions could alter the presence/absence and relative abundances of bee species in our system, bringing about substantial assemblage reordering. Given that our sampling was confined to three sites at the Sevilleta NWR, our findings represent a conservative estimate of how bee assemblages may differ among ecosystem types and thus be influenced by state transitions on a broader scale.

Our data suggest that the most probable state transitions in our southwestern U.S. drylands – shrub encroachment into Desert grassland and Desert grassland encroachment into Plains grassland^49,62^ – could substantially reshape bee communities at the Sevilleta NWR. For the grass-to-shrub transition, our results suggest that, on average over the season, total bee abundance will decrease while richness, Shannon diversity, and Pielou’s evenness will increase. In contrast, our findings predict that richness, Shannon diversity, and evenness will decrease while total abundance will remain relatively unchanged if Desert grassland replaces Plains grassland. The simultaneous occurrence of these state transitions could therefore substantially alter the distribution of bees and their ecosystem services across the landscape.

A number of factors complicate accurately predicting the outcomes of ecosystem state transitions for bees. First, bees may alter their foraging patterns based on floral resource availability^63^, and could respond to shifting vegetation composition by foraging for greater distances if floral resources are scarce in a particular location^64^. However, foraging range can differ greatly among bee species based on body size^65^, and the energetic costs of longer foraging distances may be high^66^. These factors could mediate the consequences of ecosystem state transitions for bee assemblage composition in ways that merit further research, as they have been little-examined^63^. Our finding that particular bee species were indicators of different months in different ecosystem types suggests that bees in our system may shift their foraging locations based on floral availability (though they could alternatively be emerging at different times of year in different ecosystems; see subsequent paragraphs), highlighting the importance of examining foraging dynamics in the future.

Second, state transitions may create positive feedbacks that accelerate their pace and influence bee responses^1^. Creosote bush expansion, which is limited by minimum nighttime temperature, is aided by a feedback in which a creosote individual creates a warmer microclimate around itself, which can buffer it from low temperatures, in turn creating conditions favorable to further creosote establishment^48,67^. This accelerated temperature increase could influence the relative dominances of bee species, which may differ in their temperature responses. For instance, in one *Osmia* species, increased temperature during larval development caused decreased prepupal weight and increased adult mortality^68^, and in another species it increased the frequency of 1-year rather than 2-year lifecycles^39^. If accelerated bee assemblage shifts result in pollen limitation for plants already threatened by shrub encroachment in our system, the feedback could be enhanced, further increasing the pace of encroachment.

Third, our findings may be biased by our use of passive bee trapping methods. For instance, traps are known to catch large numbers of Halictidae^50^; indeed, the two most abundant bees in our study were halictids (*Lasioglossum semicaeruleum* and *Agapostemon angelicus*). However, we captured >300 species representing 56 genera and 6 families in our study, including many pollen specialist bee species, which we caught in relatively high numbers. While absolute abundance estimates may be skewed, our methods nonetheless allow comparison of bee assemblage differences among sampling sites and seasons, as well as relative activity levels among individual bee species.

Finally, bee populations and communities can be remarkably resilient to environmental change, and their abundances are known to fluctuate substantially across space and time^40–42^. Thus, despite observed patterns related to bee assemblage differences among ecosystem types, state transitions could influence bees positively, negatively, or not at all. However, our results suggest the potential for transitions to alter communities substantially, and contribute to a number of global studies suggesting how state transitions may influence drylands arthropod communities. For instance, in the Chihuahuan Desert, ant species composition varied with mesquite (*Prosopis glandulosa*) encroachment level, but richness and abundance did not^69^. At the Sevilleta NWR, grasshopper assemblage similarity decreased with elevational variation in shrub and *Bouteloua* sp. grass cover^70^. Similarly, at our study sites, the Desert grassland and shrubland had distinct ground-dwelling arthropod assemblages, with higher abundance but similar richness in the grassland relative to the shrubland, as we found^14^. These findings together suggest that Chihuahuan Desert state transitions may similarly influence abundance and diversity patterns in several arthropod groups. However, global evidence suggests that shrub encroachment can differentially affect arthropod taxa^3^. For instance, shrub-encroached pastures in Spain had higher pollinator richness but fewer pollinator visits to forbs relative to shrub-absent sites^32^, contrasting with our finding of little difference in bee richness between shrub- and grass-dominated sites. In the Kalahari Desert, shrub encroachment corresponded with greater abundances of some ground-dwelling arthropod groups but declines in others^71^. Our results thus add to understanding of how widely-occurring ecosystem transitions may affect arthropods differentially across space.

Our work also bolsters evidence from human-altered landscapes about how bee assemblages vary at a landscape scale. For instance, as in our study, bee abundance but not richness changed with land use intensity in tropical agroecosystems for solitary bees, which comprised the majority of our dataset^29^. In contrast, a different study found shifts in bee species composition but not abundance or diversity among forest fragments^30^. Other studies have documented strong differences in both abundance and richness of bees among habitat types^33,72,73^. These findings highlight the importance of separately examining trends within particular ecosystem and land-use types to comprehensively predict future patterns.

### Shifting seasonality: phenology of bees

Month-to-month differences in bee species composition were an order of magnitude stronger than ecosystem differences in our study. This finding suggests potential susceptibility of bees in all ecosystem types to climate change-induced phenological shifts^36^. In particular, climate models for the southwestern U.S. predict less precipitation in July and August, and more in September and October, resulting in an extended period of aridity between spring rains and the start of the summer monsoon^74^. Evidence suggests that desert bees, most of which nest underground, frequently cue on precipitation for their emergence^37,38^. Under altered monsoon precipitation timing, bees that currently emerge in July or August might shift their emergence to September and October, leading to higher levels of bee abundance and richness at all sites during these months. These differences could be particularly pronounced in the Chihuahuan Desert shrubland ecosystem, for which bee abundance, Shannon *H’*, richness, and Pielou’s *J* (evenness) were all highest in July or August. The Chihuahuan Desert grassland could also be particularly susceptible to altered dynamics, given that it had the strongest seasonal turnover. Substantial assemblage reordering among months could occur if different bee species shift their phenological timing to different degrees, which could have landscape-level consequences given that the Chihuahuan Desert ecosystem types are expected to expand in the future^46–48^. In addition, for social bees that are active throughout the growing season, such as those in the family Halictidae^75^, loss of floral resources due to midsummer aridity could cause abundance declines or colony death. Predicting the consequences of ecosystem state transitions in our system will thus require considering bee assemblage seasonality in ecosystem types that are expanding versus contracting.

Regional climate predictions for the southwestern U.S. are dire – the probability of decadal droughts is nearly 100% by the end of the century^76^. Such droughts could differentially affect bees with differing phenologies and life history strategies, and could lead to bee assemblage reordering. For instance, many desert bees can remain in diapause for one year or more, emerging when conditions are favorable^37,38^. In one of the few studies on the topic, fewer bees emerged during a drought year compared to the previous and following years in the northwestern Chihuahuan Desert^38^. A greater proportion of specialist than generalist bees remained in diapause, and the specialists that emerged were those whose host plants bloom under low precipitation conditions. For *Larrea tridentata*, which requires precipitation to bloom^77^, few specialist bees emerged during the drought, suggesting that these specialists time their emergence with their host plant^38^. Differences among ecosystem types in their dominant flowering plant species, their associated specialist versus generalist bee species, and their seasonality could therefore lead to strong bee assemblage divergence among them as dominant bee species in each ecosystem respond differentially to increased drought and shifted precipitation timing, with landscape-level bee assemblage changes occurring as some ecosystems expand and others contract. Future analyses will explore connections between bee abundance, diversity, and composition and individual aspects of climate change over our time series.

Our findings of strong bee assemblage seasonality are consistent with work indicating high temporal turnover in plant-pollinator interactions in subalpine and alpine communities over the course of the growing season^41,78^. Seasonal variation in plant-pollinator networks has also been documented in agricultural landscapes^44^. While our study was not designed to examine plant-pollinator interactions, our results set the stage for considering how plant-pollinator networks could be altered by local ecosystem state transitions and climate-induced phenological shifts of bee species.

### Bee species driving among-ecosystem and within-year trends

In our dataset, common rather than rare bee species drove the trends in abundance, diversity, and composition that we observed over both space and time. Considering these species’ ecologies may be particularly important for predicting the consequences of ecosystem state transitions at the Sevilleta NWR, and some species may portend change in the bee assemblage as a whole^79^. Among the most abundant bees in our dataset, three species (*Agapostemon angelicus, Lasioglossum semicaeruleum*, and *Melissodes tristis*) were broad generalists that collect pollen from plants of many families^38,80^, suggesting that plants visited by these and other bees could be buffered to a certain extent if there are future bee declines. However, specialist bees were also among the most abundant: *Diadasia rinconis* is a specialist on Cactaceae^81^, *Anthophora affabilis* is a generalist with a strong preference for *Astragalus*, and *Perdita larreae* is a narrow specialist on *L. tridentata^23,82^*. The consequences of ecosystem state transitions for these bee species may thus depend on shifts in their host plants. For instance, expansion of *L. tridentata* could benefit populations of *P. larreae* and other creosote bush specialist bees in our system, and possibly lead to stronger competitive dynamics among creosote specialists and generalists under future conditions.

Our study identified bee species as indicators of each ecosystem type; monitoring these species, with particular attention to their life histories and level of dependence on particular plant hosts, could help to illuminate the community-level consequences of ecosystem state transitions at the Sevilleta NWR. The Chihuahuan Desert shrubland had more indicator species than the other two ecosystem types, suggesting that its future expansion could bring about distinctive assemblage shifts. The strongest indicators of the shrubland included *Perdita larreae*, which specializes on *L. tridentata*, and *P. diversa*, which specializes on *Tiquilia* spp., plants largely restricted to the shrubland site. Abundance increases of these bee species could thus signal effects of shrubland expansion on pollinator communities. Similarly, in the Plains grassland, one indicator species *(Colletes scopiventer)* specializes on *Chamaesaracha* spp., which are present in all three ecosystems but are most abundant in the Plains grassland. Future decreases in the abundance of *C. scopiventer* could signal community-level shifts accompanying the declining dominance of the Plains grassland. Specialist bee species were only indicators of ecosystem types that contained their host plants, suggesting their general utility for considering the consequences of vegetation change. However, the remaining indicator species of the two grassland ecosystems were broad generalists. Factors other than plant community composition, such as nesting habitat preferences, interspecific competitive dynamics, or floral preferences may thus underlie their restriction to particular sites, and they may be relatively less susceptible to climate-induced plant community shifts. This could also be the case for species including *Macrotera portalis*, an indicator of the shrubland but a specialist on the genus *Sphaeralcea*^83^, which is common in all three ecosystem types, and for generalist bee species that were indicators of the shrubland. The spatial distribution of suitable nesting habitat also merits future consideration in that specialist bees could be negatively affected if their plant hosts shift their ranges away from potential nesting sites.

In addition, cleptoparasitic bees were among our identified indicator species. These included one indicator of the Chihuahuan Desert shrubland (*Neolarra vigilans*) and three of the Plains grassland (*Melecta alexanderi* and two *Sphecodes* species). Not surprisingly, in the cases of *Neolarra* and *Melecta*, their bee hosts, *Perdita* and *Anthophora*, were also among indicators of the same ecosystem types, and cleptoparasitic bees were never indicators of sites where their hosts were absent. Cleptoparasitic bees may be particularly good indicators of environmental change, as they are relatively diverse and can be among the first bee functional groups to respond to disturbance^84^. Monitoring the abundances of these species could indicate shifts in bee assemblage dynamics as ecosystem state transitions occur. Cleptoparasitic species were also amongst indicators of particular months, frequently in tandem with their candidate host bees, and may thus be useful for tracking phenological responses to environmental change^84^. For example, both *Melecta alexanderi* and *M. bohartorum* were indicators of the shrubland site in March, but were indicators of both the Desert and Plains grassland sites in April, suggesting the possibility of altered emergence timing under the warmer microclimate conditions of the shrubland^48,67^, and thus susceptibility to phenological shifts in response to increasing temperature.

Our identification of indicator species may be biased by our sampling methods; species identified as indicators may be present at a site due to localized distributional fluctuations rather than habitat requirements, and species caught only at one site may be present but undetected at other sites. Our analyses nonetheless identify candidate species that may be particularly influenced by future state transitions and that merit additional consideration.

We also identified bee species that were characteristic of particular times of year across ecosystem types in our system. Monitoring these species could enable the detection of broad, cross-site phenological shifts that may occur in the future. For instance, *Osmia, Anthophora*, and *Diadasia* species may be monitored to consider shifting pre-monsoon bee phenology, and *Perdita* species may be used to study shifts in emergence timed with monsoon rains. Future publications using these data will investigate inter-annual bee assemblage differences and relationships with climate variables.

Finally, we identified bee species that were characteristic of particular months only in specific ecosystems at the Sevilleta NWR. Among these, certain specialist bee species may be candidates for detecting important phenological shifts within ecosystems, identifying phenological differences among ecosystems, and tracking how specialists versus generalists respond to climate shifts. For instance, March in the Chihuahuan Desert ecosystems had indicator species that likely specialize on Fabaceae *(Ashmeadiella erema* and *A. rubrella*), and April in the Desert and Plains grassland sites had a specialist on Brassicaceae (*Dufourea pulchricornis*). In the Chihuahuan Desert ecosystem types, specialists of creosote bush were characteristic of May, corresponding with creosote’s spring bloom^77^, June had Cactaceae specialists (*Diadasia* sp.) timed with that family’s bloom^81^, and July had specialists on Asteraceae *(Perdita ignota ignota, P. callicerata, P. fallax*, and *P. albovittata*) and *Tiquilia* sp. (*P. diversa*). In contrast, specialists on Asteraceae (*Melissodes coreopsis, P. ignota ignota*, and *P. callicerata*) were indicators of August in the Plains grassland, suggesting a shift in the importance of particular floral resources and/or differing phenological patterns among ecosystem types in our system. In August, the Chihuahuan Desert sites were characterized by numerous creosote specialists, as documented in several studies^23,38^, including *Hesperapis larreae, P. semicaerulea*, and *P. larreae*. These species may be candidates for examining how delayed monsoon influences bee phenology. The *Sphaeralcea* specialist *Macrotera portalis*^83^ was an August indicator in the shrubland, but other *Sphaeralcea* specialists were characteristic of September in both Chihuahuan Desert ecotypes; perhaps competitive dynamics were responsible for this difference. These examples illustrate the suite of factors that could be important to consider in order to predict bee presence and seasonality across the Sevilleta NWR landscape.

### Summary

Our analysis of 13 years of bee assemblage data spanning 302 species suggests that future dryland ecosystem state transitions, by themselves, may alter bee species’ relative abundances and presence/absence in our system. Strong bee assemblage seasonal turnover, particularly in ecosystems predicted to expand, indicates the potential for bee phenological shifts to accompany state transitions, potentially reordering communities. Our results indicate that predicting the consequences of global change for bee assemblages will require accounting for both within-year and among-ecosystem variation.

## Data availability

The data generated and analyzed during the current study are available in the Environmental Data Initiative (EDI) Data Portal (https://doi.org/10.6073/pasta/efae7b928d60e0caa3ac1268832f268f).

## Acknowledgements

Funding was provided by the NSF Long Term Ecological Research program (DEB-1655499) and the University of New Mexico Department of Biology. We thank David Lightfoot, Olivia Messinger Carril, and Jade McLaughlin for their contributions to this work.

## Author contributions

M.R.K. assisted with bee specimen collection, analyzed the data, and wrote the manuscript. K.W.W. designed the study and completed the majority of specimen collection and identification. J.B. assisted with specimen collection and identification. T.L.G. identified specimens and provided taxonomic expertise. J.A.R. and K.D.W. contributed to manuscript conceptual framing, data analysis, and writing. All authors contributed to revising the manuscript.

## References

1. D’Odorico, P., Bhattachan, A., Davis, K. F., Ravi, S. & Runyan, C. W. Global desertification: drivers and feedbacks. Adv. Water Resour. 51, 326–344 (2013).

2. Bestelmeyer, B. T. et al. Desertification, land use, and the transformation of global drylands. Front. Ecol. Environ. 13, 28–36 (2015).

3. Eldridge, D. J. et al. Impacts of shrub encroachment on ecosystem structure and functioning: towards a global synthesis. Ecol. Lett. 14, 709–722 (2011).

4. Allen, C. D., Breshears, D. D. & McDowell, N. G. On underestimation of global vulnerability to tree mortality and forest die-off from hotter drought in the Anthropocene. Ecosphere 6, art129 (2015).

5. Sala, O. E. & Maestre, F. T. Grass-woodland transitions: determinants and consequences for ecosystem functioning and provisioning of services. J. Ecol. 102, 1357–1362 (2014).

6. Biederman, J. A. et al. Terrestrial carbon balance in a drier world: the effects of water availability in southwestern North America. Glob. Change Biol. 22, 1867–1879 (2016).

7. Bestelmeyer, B. T. et al. The grassland–shrubland regime shift in the southwestern United States: misconceptions and their implications for management. Bioscience 68, 678–690 (2018).

8. Anderson-Teixeira, K. J., Delong, J. P., Fox, A. M., Brese, D. A. & Litvak, M. E. Differential responses of production and respiration to temperature and moisture drive the carbon balance across a climatic gradient in New Mexico. Glob. Change Biol. 17, 410–424 (2011).

9. Petrie, M. D., Collins, S. L., Swann, A. M., Ford, P. L. & Litvak, M. E. Grassland to shrubland state transitions enhance carbon sequestration in the northern Chihuahuan Desert. Glob. Change Biol. 21, 1226–1235 (2015).

10. Turnbull, L., Wainwright, J. & Brazier, R. E. Nitrogen and phosphorus dynamics during runoff events over a transition from grassland to shrubland in the south-western United States. Hydrol. Process. 25, 1–17 (2011).

11. Wang, G. et al. Post-fire redistribution of soil carbon and nitrogen at a grassland–shrubland ecotone. Ecosystems 22, 174–188 (2019).

12. Ratajczak, Z. et al. Changes in spatial variance during a grassland to shrubland state transition. J. Ecol. 105, 750–760 (2017).

13. Sanchez, B. C. & Parmenter, R. R. Patterns of shrub-dwelling arthropod diversity across a desert shrubland-grassland ecotone: a test of island biogeographic theory. J. Arid Environ. 50, 247–265 (2002).

14. Lightfoot, D. C., Brantley, S. L. & Allen, C. D. Geographic patterns of ground-dwelling arthropods across an ecoregional transition in the North American Southwest. West. North Am. Nat. 68, 83–102 (2008).

15. Pravalie, R. Drylands extent and environmental issues: a global approach. Earth-Sci. Rev. 161, 259–278 (2016).

16. UN EMG. Global drylands: a UN system-wide response. Environ. Manag. Group U. N. Geneva http://www.unccd.int/Lists/SiteDocumentLibrary/Publications/Global_Drylands_Full_Report.pdf (2011).

17. Kovács-Hostyánszki, A. et al. Earthworms, spiders and bees as indicators of habitat quality and management in a low-input farming region—a whole farm approach. Ecol. Indic. 33, 111–120 (2013).

18. Gonçalves, R. B., Sydney, N. V., Oliveira, P. S. & Artmann, N. O. Bee and wasp responses to a fragmented landscape in southern Brazil. J. Insect Conserv. 18, 1193–1201 (2014).

19. Klein, A.-M. et al. Importance of pollinators in changing landscapes for world crops. Proc. R. Soc. B Biol. Sci. 274, 303–313 (2007).

20. Ollerton, J., Winfree, R. & Tarrant, S. How many flowering plants are pollinated by animals? Oikos 120, 321–326 (2011).

21. Michener, C. D. The Bees of the World. (Johns Hopkins University Press, 2007).

22. Minckley, R. L. & Ascher, J. S. Preliminary survey of bee (Hymenoptera: Anthophila) richness in the northwestern Chihuahuan Desert in Merging Science and Management in a Rapidly Changing World: Biodiversity and Management of the Madrean Archipelago III and 7th Conference on Research and Resource Management in the Southwestern Deserts, Proceedings RMRS-P-67 (eds. Gottfried, G. J., Ffolliott, P. F., Gebow, B. S., Eskew, L. G. & Collins, L. C.) 138–143 (U.S. Department of Agriculture, Forest Service, Rocky Mountain Research Station, 2013).

23. Minckley, R. L., Cane, J. H. & Kervin, L. Origins and ecological consequences of pollen specialization among desert bees. Proc. R. Soc. Lond. B Biol. Sci. 267, 265–271 (2000).

24. Turner, R. M., Bowers, J. E. & Brugess, T. L. Sonoran Desert Plants: An Ecological Atlas. (The University of Arizona Press, 2005).

25. Minckley, R. L., Cane, J. H., Kervin, L. & Roulston, T. H. Spatial predictability and resource specialization of bees (Hymenoptera: Apoidea) at a superabundant, widespread resource. Biol. J. Linn. Soc. 67, 119–147 (1999).

26. Simpson, B. & Neff, J. Pollination ecology in the Southwest. Aliso 11, 417–440 (1987).

27. Memmott, J., Craze, P. G., Waser, N. M. & Price, M. V. Global warming and the disruption of plant-pollinator interactions. Ecol. Lett. 10, 710–717 (2007).

28. Kaiser-Bunbury, C. N., Muff, S., Memmott, J., Müller, C. B. & Caflisch, A. The robustness of pollination networks to the loss of species and interactions: a quantitative approach incorporating pollinator behaviour. Ecol. Lett. 13, 442–452 (2010).

29. Klein, A.-M., Steffan-Dewenter, I., Buchori, D. & Tscharntke, T. Effects of land-use intensity in tropical agroforestry systems on coffee flower-visiting and trap-nesting bees and wasps. Conserv. Biol. 16, 1003–1014 (2002).

30. Brosi, B. J., Daily, G. C., Shih, T. M., Oviedo, F. & Durán, G. The effects of forest fragmentation on bee communities in tropical countryside. J. Appl. Ecol. 45, 773–783 (2008).

31. Banaszak-Cibicka, W. & Żmihorski, M. Wild bees along an urban gradient: winners and losers. J. Insect Conserv. 16, 331–343 (2012).

32. Lara-Romero, C., García, C., Morente-López, J. & Iriondo, J. M. Direct and indirect effects of shrub encroachment on alpine grasslands mediated by plant-flower visitor interactions. Funct. Ecol. 30, 1521–1530 (2016).

33. Minckley, R. Faunal composition and species richness differences of bees (Hymenoptera: Apiformes) from two north American regions. Apidologie 39, 176–188 (2008).

34. Torné-Noguera, A. et al. Determinants of spatial distribution in a bee community: nesting resources, flower resources, and body size. PLoS ONE 9, e97255 (2014).

35. Ruttan, A., Filazzola, A. & Lortie, C. J. Shrub-annual facilitation complexes mediate insect community structure in arid environments. J. Arid Environ. 134, 1–9 (2016).

36. Renner, S. S. & Zohner, C. M. Climate change and phenological mismatch in trophic interactions among plants, insects, and vertebrates. Annu. Rev. Ecol. Evol. Syst. 49, 165–182 (2018).

37. Danforth, B. N. Emergence dynamics and bet hedging in a desert bee, *Perdita portalis*. Proc. R. Soc. Lond. B Biol. Sci. 266, 1985–1994 (1999).

38. Minckley, R. L., Roulston, T. H. & Williams, N. M. Resource assurance predicts specialist and generalist bee activity in drought. Proc. R. Soc. B Biol. Sci. 280, 20122703 (2013).

39. Forrest, J. R. K., Cross, R. & CaraDonna, P. J. Two-year bee, or not two-year bee? How voltinism Is affected by temperature and season length in a high-elevation solitary bee. Am. Nat. 193, 560–574 (2019).

40. Tylianakis, J. M., Klein, A.-M. & Tscharntke, T. Spatiotemporal variation in the diversity of Hymenoptera across a tropical habitat gradient. Ecology 86, 3296–3302 (2005).

41. Simanonok, M. P. & Burkle, L. A. Partitioning interaction turnover among alpine pollination networks: spatial, temporal, and environmental patterns. Ecosphere 5, art149 (2014).

42. Rollin, O., Bretagnolle, V., Fortel, L., Guilbaud, L. & Henry, M. Habitat, spatial and temporal drivers of diversity patterns in a wild bee assemblage. Biodivers. Conserv. 24, 1195–1214 (2015).

43. Leong, M., Ponisio, L. C., Kremen, C., Thorp, R. W. & Roderick, G. K. Temporal dynamics influenced by global change: bee community phenology in urban, agricultural, and natural landscapes. Glob. Change Biol. 22, 1046–1053 (2016).

44. Tucker, E. M. & Rehan, S. M. Farming for bees: annual variation in pollinator populations across agricultural landscapes. Agric. For. Entomol. 20, 541–548 (2018).

45. Notaro, M. et al. Complex seasonal cycle of ecohydrology in the Southwest United States. J. Geophys. Res. Biogeosciences 115, G04034 (2010).

46. Peters, D. P. C. & Yao, J. Long-term experimental loss of foundation species: consequences for dynamics at ecotones across heterogeneous landscapes. Ecosphere 3, art27 (2012).

47. Collins, S. L. & Xia, Y. Long-term dynamics and hotspots of change in a desert grassland plant community. Am. Nat. 185, E30–E43 (2015).

48. He, Y., D’Odorico, P. & De Wekker, S. F. J. The role of vegetation-microclimate feedback in promoting shrub encroachment in the northern Chihuahuan desert. Glob. Change Biol. 21, 2141–2154 (2015).

49. Rudgers, J. A. et al. Climate sensitivity functions and net primary production: a framework for incorporating climate mean and variability. Ecology 99, 576–582 (2018).

50. Wilson, J. S., Griswold, T. & Messinger, O. J. Sampling bee communities (Hymenoptera: Apiformes) in a desert landscape: are pan traps sufficient? J. Kans. Entomol. Soc. 81, 288–300 (2008).

51. Cane, J. H., Minckley, R. L. & Kervin, L. J. Sampling bees (Hymenoptera: Apiformes) for pollinator community studies: pitfalls of pan-trapping. J. Kans. Entomol. Soc. 73, 225–231 (2000).

52. Baum, K. A. & Wallen, K. E. Potential bias in pan trapping as a function of floral abundance. J. Kans. Entomol. Soc. 84, 155–159 (2011).

53. Leong, J. M. & Thorp, R. W. Colour-coded sampling: the pan trap colour preferences of oligolectic and nonoligolectic bees associated with a vernal pool plant. Ecol. Entomol. 24, 329–335 (1999).

54. Wickham, H. Reshaping data with the reshape package. J. Stat. Softw. 21, 10.18637/jss.v021.i12 (2007).

55. R Core Team. R: A Language and Environment for Statistical Computing. (R Foundation for Statistical Computing, 2017).

56. Clarke, K. R. & Gorley, R. N. Primer Version 6.1.10 User Manual and Tutorial. (Primer-E, 2009).

57. Hallett, L. et al. codyn: community dynamics metrics, R package version 2.0.2. 10.5063/F1N877Z6 (2019).

58. Roberts, D. W. labdsv: ordination and multivariate analysis for ecology. https://CRAN.R-project.org/package=labdsv (2016).

59. Oksanen, J. et al. vegan: community ecology package, version 2.2-1. http://CRAN.R-project.org/package=vegan (2015).

60. Bates, D., Mächler, M., Bolker, B. & Walker, S. Fitting linear mixed-effects models using lme4. J. Stat. Softw. 67, 1–48 (2015).

61. Lenth, R. V., Singmann, H., Love, J., Buerkner, P. & Herve, M. emmeans: estimated marginal means, aka least-squares means. https://cran.r-project.org/web/packages/emmeans/index.html (2018).

62. Collins, S. L. et al. A multiscale, hierarchical model of pulse dynamics in arid-land ecosystems. Annu. Rev. Ecol. Evol. Syst. 45, 397–419 (2014).

63. Ogilvie, J. E. & Forrest, J. R. Interactions between bee foraging and floral resource phenology shape bee populations and communities. Curr. Opin. Insect Sci. 21, 75–82 (2017).

64. Redhead, J. W. et al. Effects of habitat composition and landscape structure on worker foraging distances of five bumble bee species. Ecol. Appl. 26, 726–739 (2016).

65. Greenleaf, S. S., Williams, N. M., Winfree, R. & Kremen, C. Bee foraging ranges and their relationship to body size. Oecologia 153, 589–596 (2007).

66. Horn, J., Becher, M. A., Kennedy, P. J., Osborne, J. L. & Grimm, V. Multiple stressors: using the honeybee model BEEHAVE to explore how spatial and temporal forage stress affects colony resilience. Oikos 125, 1001–1016 (2016).

67. He, Y., D’Odorico, P., De Wekker, S. F. J., Fuentes, J. D. & Litvak, M. On the impact of shrub encroachment on microclimate conditions in the northern Chihuahuan desert. J. Geophys. Res.-Atmospheres 115, D21120 (2010).

68. Radmacher, S. & Strohm, E. Effects of constant and fluctuating temperatures on the development of the solitary bee *Osmia bicornis* (Hymenoptera: Megachilidae). Apidologie 42, 711–720 (2011).

69. Bestelmeyer, B. T. Does desertification diminish biodiversity? Enhancement of ant diversity by shrub invasion in south-western USA. Divers. Distrib. 11, 45–55 (2005).

70. Rominger, A. J., Miller, T. E. X. & Collins, S. L. Relative contributions of neutral and niche-based processes to the structure of a desert grassland grasshopper community. Oecologia 161, 791–800 (2009).

71. Blaum, N., Seymour, C., Rossmanith, E., Schwager, M. & Jeltsch, F. Changes in arthropod diversity along a land use driven gradient of shrub cover in savanna rangelands: identification of suitable indicators. Biodivers. Conserv. 18, 1187–1199 (2009).

72. Bates, A. J. et al. Changing bee and hoverfly pollinator assemblages along an urban-rural gradient. PLoS ONE 6, e23459 (2011).

73. Tonietto, R., Fant, J., Ascher, J., Ellis, K. & Larkin, D. A comparison of bee communities of Chicago green roofs, parks and prairies. Landsc. Urban Plan. 103, 102–108 (2011).

74. Cook, B. I. & Seager, R. The response of the North American Monsoon to increased greenhouse gas forcing. J. Geophys. Res. Atmospheres 118, 1690–1699 (2013).

75. Michener, C. D. The Social Behavior of the Bees: A Comparative Study. (Belknap Press of Harvard University Press, 1974).

76. Cook, B. I., Ault, T. R. & Smerdon, J. E. Unprecedented 21st century drought risk in the American Southwest and Central Plains. Sci. Adv. 1, e1400082 (2015).

77. Bowers, J. E. & Dimmitt, M. A. Flowering phenology of six woody plants in the Northern Sonoran Desert. Bull. Torrey Bot. Club 121, 215–229 (1994).

78. CaraDonna, P. J. et al. Interaction rewiring and the rapid turnover of plant–pollinator networks. Ecol. Lett. 20, 385–394 (2017).

79. Ellison, A. M. et al. Loss of foundation species: consequences for the structure and dynamics of forested ecosystems. Front. Ecol. Environ. 3, 479–486 (2005).

80. Schuh, R. T., Hewson-Smith, S. & Ascher, J. S. Specimen databases: a case study in entomology using web-based software. Am. Entomol. 56, 206–216 (2010).

81. Sipes, S. D. & Tepedino, V. J. Pollen-host specificity and evolutionary patterns of host switching in a clade of specialist bees (Apoidea: *Diadasia*). Biol. J. Linn. Soc. 86, 487–505 (2005).

82. Hurd, P. D. & Linsley, E. G. The principal *Larrea* bees of the southwestern United States (Hymenoptera, Apoidea). Smithson. Contrib. Zool. 1–74 (1975).

83. Danforth, B. N., Ji, S. & Ballard, L. J. Gene flow and population structure in an oligolectic desert bee, *Macrotera (Macroteropsis) portalis* (Hymenoptera: Andrenidae). J. Kans. Entomol. Soc. 76, 221–235 (2003).

84. Sheffield, C. S., Pindar, A., Packer, L. & Kevan, P. G. The potential of cleptoparasitic bees as indicator taxa for assessing bee communities. Apidologie 44, 501–510 (2013).

