## Supplementary Material for "Predicting changes in bee assemblages following state transitions at North American dryland ecotones"

### Supplementary Information

Includes:
Supplementary Tables S1, S2, S3, S4, S5, S6
Supplementary Figures S1, S2, S3, S4, S5, S6

**Supplementary Table S1.** Bee species collected during the study. In the Species column, numbers or codes are used as unique identifiers in certain cases where species have not yet been described. Author indicates the authority who described the species.

| Family | Genus | Subgenus | Species | Author | Notes |
|---|---|---|---|---|---|
| Andrenidae | *Andrena* | *Callandrena* | *ardis* | LaBerge | |
| Andrenidae | *Andrena* | *Callandrena* | *pecosana* | Cockerell | |
| Andrenidae | *Andrena* | *Micrandrena* | 2 | | New species |
| Andrenidae | *Andrena* | *Micrandrena* | *illinoiensis* | Robertson | |
| Andrenidae | *Andrena* | *Micrandrena* | *primulifrons* | Casad | |
| Andrenidae | *Andrena* | NA | 3 | | |
| Andrenidae | *Andrena* | *Plastandrena* | *prunorum* | Cockerell | |
| Andrenidae | *Andrena* | *Rhaphandrena* | *prima* | Casad | |
| Andrenidae | *Andrena* | *Tylandrena* | *jessicae* | Cockerell | |
| Andrenidae | *Andrena* | *Tylandrena* | *mesillae* | Cockerell | |
| Andrenidae | *Calliopsis* | NA | 2 | | |
| Andrenidae | *Calliopsis* | *Nomadopsis* | *meliloti* | Cockerell | |
| Andrenidae | *Calliopsis* | *Perissander* | *anomoptera* | Michener | |
| Andrenidae | *Calliopsis* | *Verbenapis* | *verbenae* | Cockerell & Porter | |
| Andrenidae | *Macrotera* | *Macroteropsis* | *latior* | (Cockerell) | |
| Andrenidae | *Macrotera* | *Macroteropsis* | *magniceps* | (Timberlake) | |
| Andrenidae | *Macrotera* | *Macroteropsis* | *portalis* | (Timberlake) | |
| Andrenidae | *Perdita* | *Cockerellia* | *albipennis heliophila* | Cresson | |
| Andrenidae | *Perdita* | *Cockerellia* | *coreopsidis kansensis* | Timberlake | |
| Andrenidae | *Perdita* | *Cockerellia* | *verbesinae* | Cockerell | |
| Andrenidae | *Perdita* | *Epimacrotera* | *diversa* | Timberlake | |
| Andrenidae | *Perdita* | *Hexaperdita* | 4 | | |
| Andrenidae | *Perdita* | *Hexaperdita* | aff. *heterotheceae* | | |
| Andrenidae | *Perdita* | *Hexaperdita* | *callicerata* | Cockerell | |
| Andrenidae | *Perdita* | *Hexaperdita* | *cara* | Timberlake | |
| Andrenidae | *Perdita* | *Hexaperdita* | *ignota ignota* | Cockerell | |
| Andrenidae | *Perdita* | NA | 10 | | New species |
| Andrenidae | *Perdita* | NA | 14 | | |
| Andrenidae | *Perdita* | NA | 15 | | |
| Andrenidae | *Perdita* | NA | 16 | | |
| Andrenidae | *Perdita* | NA | 17 | | |
| Andrenidae | *Perdita* | NA | 18 | | |
| Andrenidae | *Perdita* | NA | 19 | | |
| Andrenidae | *Perdita* | NA | 20 | | |
| Andrenidae | *Perdita* | NA | 21 | | |
| Andrenidae | *Perdita* | NA | 33 | | |
| Andrenidae | *Perdita* | *Pentaperdita* | 3 | | |
| Andrenidae | *Perdita* | *Pentaperdita* | aff. *albovittata* | | |
| Andrenidae | *Perdita* | *Pentaperdita* | *albovittata* | Cockerell | |
| Andrenidae | *Perdita* | *Pentaperdita* | *bradleyana* | Timberlake | |
| Andrenidae | *Perdita* | *Perdita* | 2 | | Sphaeralceae group |
| Andrenidae | *Perdita* | *Perdita* | 5 | | Sphaeralceae group |
| Andrenidae | *Perdita* | *Perdita* | 6 | | New species, Octomaculata group |
| Andrenidae | *Perdita* | *Perdita* | 11 | | New species, Octomaculata group |
| Andrenidae | *Perdita* | *Perdita* | 12 | | New species, Octomaculata group |
| Andrenidae | *Perdita* | *Perdita* | 13 | | Chamaesarachae group |
| Andrenidae | *Perdita* | *Perdita* | 31 | | New species, Octomaculata group |
| Andrenidae | *Perdita* | *Perdita* | 32 | | New species |
| Andrenidae | *Perdita* | *Perdita* | 36 | | New species |
| Andrenidae | *Perdita* | *Perdita* | 50 | | Sphaeralceae group |
| Andrenidae | *Perdita* | *Perdita* | aff. *cuspidata* | | |

| Family | Genus | Subgenus | Species | Author | Notes |
|---|---|---|---|---|---|
| Andrenidae | *Perdita* | *Perdita* | aff. *laticincta* | | |
| Andrenidae | *Perdita* | *Perdita* | aff. *missionis* | | |
| Andrenidae | *Perdita* | *Perdita* | *affinis* | Cresson | |
| Andrenidae | *Perdita* | *Perdita* | *aperta* | Timberlake | |
| Andrenidae | *Perdita* | *Perdita* | *aridella* | Timberlake | |
| Andrenidae | *Perdita* | *Perdita* | *austini* | Cockerell | |
| Andrenidae | *Perdita* | *Perdita* | *confusa* | Timberlake | |
| Andrenidae | *Perdita* | *Perdita* | *drymariae* | Timberlake | |
| Andrenidae | *Perdita* | *Perdita* | *fallax* | Cockerell | |
| Andrenidae | *Perdita* | *Perdita* | *gutierreziae* | Cockerell | |
| Andrenidae | *Perdita* | *Perdita* | *lenis* | Timberlake | |
| Andrenidae | *Perdita* | *Perdita* | *luteola* | Cockerell | |
| Andrenidae | *Perdita* | *Perdita* | *rectangulata* | Cockerell | |
| Andrenidae | *Perdita* | *Perdita* | *semicaerulea* | Cockerell | |
| Andrenidae | *Perdita* | *Perdita* | *sphaeralceae alticola* | Cockerell | |
| Andrenidae | *Perdita* | *Perdita* | *trinotata* | Timberlake | |
| Andrenidae | *Perdita* | *Perditella* | *larreae* | Cockerell | |
| Andrenidae | *Perdita* | *Perditella* | *marcialis* | Cockerell | |
| Andrenidae | *Protandrena* | *Heterosarus* | 1 | | |
| Andrenidae | *Protandrena* | *Heterosarus* | 2 | | |
| Andrenidae | *Protandrena* | *Heterosarus* | 3 | | |
| Andrenidae | *Protoxaea* | NA | *gloriosa* | (Fox) | |
| Andrenidae | *Pseudopanurgus* | NA | *fraterculus fraterculus* | (Cockerell) | |
| Apidae | *Anthophora* | *Anthophoroides* | *californica* | Cresson | |
| Apidae | *Anthophora* | *Heliophila* | *petrophila* | Cockerell | |
| Apidae | *Anthophora* | *Heliophila* | *phenax* | (Cockerell) | |
| Apidae | *Anthophora* | *Lophanthophora* | *affabilis* | Cresson | |
| Apidae | *Anthophora* | *Lophanthophora* | *coptognatha* | Timberlake | |
| Apidae | *Anthophora* | *Lophanthophora* | *neglecta* | Timberlake & Cockerell | |
| Apidae | *Anthophora* | *Lophanthophora* | *porterae* | Cockerell | |
| Apidae | *Anthophora* | *Mystacanthophora* | *montana* | Cresson | |
| Apidae | *Anthophora* | *Mystacanthophora* | *urbana* | Cresson | |
| Apidae | *Anthophora* | NA | n. sp. | | Author: Brooks, manuscript in prep |
| Apidae | *Anthophora* | *Paramegilla* | *centriformis* | Cresson | |
| Apidae | *Anthophora* | *Pyganthophora* | *lesquerellae* | (Cockerell) | |
| Apidae | *Anthophorula* | *Anthophorisca* | *pygmaea* | (Cresson) | |
| Apidae | *Anthophorula* | *Anthophorula* | *compactula* | Cockerell | |
| Apidae | *Anthophorula* | *Anthophorula* | *completa* | (Cockerell) | |
| Apidae | *Apis* | *Apis* | *mellifera* | Linnaeus | |
| Apidae | *Bombus* | *Thoracobombus* | *pensylvanicus* | (DeGeer) | |
| Apidae | *Centris* | *Paracentris* | *atripes* | Mocsary | |
| Apidae | *Centris* | *Paracentris* | *caesalpiniae* | Cockerell | |
| Apidae | *Centris* | *paracentris* | *lanosa* | Cresson | |
| Apidae | *Centris* | *paracentris* | *zacateca* | Snelling | |
| Apidae | *Ceratina* | *Zadontomerus* | *nanula* | Cockerell | |
| Apidae | *Diadasia* | NA | *australis* | (Cresson) | |
| Apidae | *Diadasia* | NA | *diminuta* | (Cresson) | |
| Apidae | *Diadasia* | NA | *enavata* | (Cresson) | |
| Apidae | *Diadasia* | NA | *megamorpha* | Cockerell | |
| Apidae | *Diadasia* | NA | *ochracea* | (Cockerell) | |
| Apidae | *Diadasia* | NA | *rinconis* | Cockerell | |
| Apidae | *Epeolus* | NA | aff. *americanus* | | |
| Apidae | *Epeolus* | NA | *brumleyi* | Onuferko | |
| Apidae | *Epeolus* | NA | *compactus* | Cresson | |
| Apidae | *Epeolus* | NA | *mesillae* | (Cockerell) | |
| Apidae | *Epeolus* | NA | *minimus* | (Robertson) | |
| Apidae | *Ericrocis* | NA | *lata* | (Cresson) | |
| Apidae | *Eucera* | *Peponapis* | *pruinosa* | (Say) | |
| Apidae | *Eucera* | *Synhalonia* | aff. *albescens* | | |

| Family | Genus | Subgenus | Species | Author | Notes |
|---|---|---|---|---|---|
| Apidae | *Eucera* | *Synhalonia* | *illinoensis* | (Robertson) | |
| Apidae | *Eucera* | *Synhalonia* | *lycii* | (Cockerell) | |
| Apidae | *Eucera* | *Synhalonia* | *territella* | (Cockerell) | |
| Apidae | *Eucera* | *Tetraloniella* | *albata* | (Cresson) | |
| Apidae | *Eucera* | *Tetraloniella* | *eriocarpi* | (Cockerell) | |
| Apidae | *Eucera* | *Tetraloniella* | *silacea?* | LaBerge | Unconfirmed identification |
| Apidae | *Exomalopsis* | *Phanomalopsis* | *solani* | Cockerell | |
| Apidae | *Habropoda* | NA | *morrisoni* | (Cresson) | |
| Apidae | *Habropoda* | NA | *pallida* | (Timberlake) | |
| Apidae | *Holcopasites* | NA | *calliopsidis* | (Linsley) | |
| Apidae | *Holcopasites* | NA | *pulchellus* | (Cresson) | |
| Apidae | *Martinapis* | NA | *luteicornis* | (Cockerell) | |
| Apidae | *Melecta* | *Melecta* | *alexanderi* | Griswold & Parker | |
| Apidae | *Melecta* | *Melecta* | *bohartorum* | Linsley | |
| Apidae | *Melecta* | *Melecta* | *pacifica* | Cresson | |
| Apidae | *Melissodes* | *Callimelissodes* | *coloradensis* | Cresson | |
| Apidae | *Melissodes* | *Eumelissodes* | *agilis* | Cresson | |
| Apidae | *Melissodes* | *Eumelissodes* | *bimatris* | LaBerge | |
| Apidae | *Melissodes* | *Eumelissodes* | *coreopsis* | Robertson | |
| Apidae | *Melissodes* | *Eumelissodes* | *fasciatella* | LaBerge | |
| Apidae | *Melissodes* | *Eumelissodes* | *menuachus* | LaBerge | |
| Apidae | *Melissodes* | *Eumelissodes* | *montanus* | Cresson | |
| Apidae | *Melissodes* | *Eumelissodes* | *pallidisignatus* | Cockerell | |
| Apidae | *Melissodes* | *Eumelissodes* | *perpolitus* | LaBerge | |
| Apidae | *Melissodes* | *Eumelissodes* | *semilupinus* | Cockerell | |
| Apidae | *Melissodes* | *Eumelissodes* | *snowii* | Cresson | |
| Apidae | *Melissodes* | *Eumelissodes* | *subagilis* | Cockerell | |
| Apidae | *Melissodes* | *Eumelissodes* | *submenuacha* | Cockerell | |
| Apidae | *Melissodes* | *Eumelissodes* | *tristis* | Cockerell | |
| Apidae | *Melissodes* | *Eumelissodes* | *verbesinarum* | Cockerell | |
| Apidae | *Melissodes* | *Melissodes* | *communis communis* | Cresson | |
| Apidae | *Melissodes* | *Melissodes* | *gilensis* | Cockerell | |
| Apidae | *Melissodes* | *Melissodes* | *paroselae* | Cockerell | |
| Apidae | *Melissodes* | *Melissodes* | *thelypodii thelypodii* | Cockerell | |
| Apidae | *Neolarra* | NA | *vigilans* | (Cockerell) | |
| Apidae | *Nomada* | NA | spp. | | All Nomada were grouped together, as they could not be identified to species. |
| Apidae | *Svastra* | *Anthedonia* | *compta* | (Cresson) | |
| Apidae | *Svastra* | *Epimelissodes* | *helianthelli* | (Cockerell) | |
| Apidae | *Svastra* | *Epimelissodes* | *machaerantherae* | (Cockerell) | |
| Apidae | *Svastra* | *Epimelissodes* | *obliqua* | (Say) | |
| Apidae | *Svastra* | *Epimelissodes* | *sabinensis* | (Cockerell) | |
| Apidae | *Townsendiella* | NA | *pulchra* | Crawford | |
| Apidae | *Triepeolus* | NA | 5 | | |
| Apidae | *Triepeolus* | NA | 9 | | |
| Apidae | *Triepeolus* | NA | *grandis* | (Friese) | |
| Apidae | *Triepeolus* | NA | *norae* | Cockerell | |
| Apidae | *Xeromelecta* | NA | *interrupta* | (Cresson) | |
| Apidae | *Xylocopa* | NA | *californica* | Cresson | |
| Apidae | *Zacosmia* | NA | *maculata* | (Cresson) | |
| Colletidae | *Caupolicana* | NA | *yarrowi* | (Cresson) | |
| Colletidae | *Colletes* | NA | *birkmanni* | Swenk | |
| Colletidae | *Colletes* | NA | *clypeonitens* | Swenk | |
| Colletidae | *Colletes* | NA | *compactus* | Cresson | |
| Colletidae | *Colletes* | NA | *covilleae* | Timberlake | |
| Colletidae | *Colletes* | NA | *phaceliae* | Cockerell | |

| Family | Genus | Subgenus | Species | Author | Notes |
|---|---|---|---|---|---|
| Colletidae | *Colletes* | NA | *salicicola* | Cockerell | |
| Colletidae | *Colletes* | NA | *scopiventer* | Swenk | |
| Colletidae | *Colletes* | NA | *sphaeralceae* | Timberlake | |
| Colletidae | *Colletes* | NA | *wootoni* | Cockerell | |
| Colletidae | *Hylaeus* | *Paraprosopis* | 1 | | |
| Colletidae | *Hylaeus* | *Paraprosopis* | 2 | | |
| Colletidae | *Hylaeus* | *Paraprosopis* | *asininus* | (Cockerell & Casad) | |
| Colletidae | *Hylaeus* | *Prosopis* | *episcopelis coquilletti* | (Cockerell) | |
| Halictidae | *Agapostemon* | NA | *angelicus* | Cockerell | |
| Halictidae | *Agapostemon* | NA | *melliventris* | Cresson | |
| Halictidae | *Agapostemon* | NA | *obliquus* | (Provancher) | |
| Halictidae | *Agapostemon* | NA | *splendens* | (Lepeletier) | |
| Halictidae | *Agapostemon* | NA | *virescens* | (Fabricius) | |
| Halictidae | *Augochlorella* | NA | *aurata* | (Smith) | |
| Halictidae | *Conanthalictus* | NA | *conanthi* | (Cockerell) | |
| Halictidae | *Dufourea* | NA | *pulchricornis* | (Cockerell) | |
| Halictidae | *Dufourea* | NA | *vernalis* | Timberlake | |
| Halictidae | *Halictus* | NA | *ligatus* | Say | |
| Halictidae | *Halictus* | NA | *tripartitus* | Cockerell | |
| Halictidae | *Lasioglossum* | *Afrodialictus* | *lampronotum* | (Cameron) | |
| Halictidae | *Lasioglossum* | *Dialictus* | 2 | | |
| Halictidae | *Lasioglossum* | *Dialictus* | 4 | | |
| Halictidae | *Lasioglossum* | *Dialictus* | 5 | | |
| Halictidae | *Lasioglossum* | *Dialictus* | 7 | | |
| Halictidae | *Lasioglossum* | *Dialictus* | 8 | | |
| Halictidae | *Lasioglossum* | *Dialictus* | 11 | | |
| Halictidae | *Lasioglossum* | *Dialictus* | 12 | | |
| Halictidae | *Lasioglossum* | *Dialictus* | 13 | | |
| Halictidae | *Lasioglossum* | *Dialictus* | 19 | | |
| Halictidae | *Lasioglossum* | *Dialictus* | 20 | | |
| Halictidae | *Lasioglossum* | *Dialictus* | 21 | | |
| Halictidae | *Lasioglossum* | *Dialictus* | 30 | | |
| Halictidae | *Lasioglossum* | *Dialictus* | 32 | | |
| Halictidae | *Lasioglossum* | *Dialictus* | 33 | | |
| Halictidae | *Lasioglossum* | *Dialictus* | 34 | | |
| Halictidae | *Lasioglossum* | *Dialictus* | 35 | | |
| Halictidae | *Lasioglossum* | *Dialictus* | 36 | | |
| Halictidae | *Lasioglossum* | *Dialictus* | 37 | | |
| Halictidae | *Lasioglossum* | *Dialictus* | 38 | | |
| Halictidae | *Lasioglossum* | *Dialictus* | 39 | | |
| Halictidae | *Lasioglossum* | *Dialictus* | 40 | | |
| Halictidae | *Lasioglossum* | *Dialictus* | 42 | | |
| Halictidae | *Lasioglossum* | *Dialictus* | 46 | | |
| Halictidae | *Lasioglossum* | *Dialictus* | 47 | | |
| Halictidae | *Lasioglossum* | *Dialictus* | 48 | | |
| Halictidae | *Lasioglossum* | *Dialictus* | aff. *pervarum* | | |
| Halictidae | *Lasioglossum* | *Dialictus* | cf. *coactum* | | |
| Halictidae | *Lasioglossum* | *Dialictus* | *comulum* | Michener | |
| Halictidae | *Lasioglossum* | *Dialictus* | *eophilum* | (Ellis) | |
| Halictidae | *Lasioglossum* | *Dialictus* | *hudsoniellum* | (Cockerell) | |
| Halictidae | *Lasioglossum* | *Dialictus* | *microlepoides* | (Ellis) | |
| Halictidae | *Lasioglossum* | *Dialictus* | n. sp. | | New species |
| Halictidae | *Lasioglossum* | *Dialictus* | *semicaeruleum* | (Cockerell) | |
| Halictidae | *Lasioglossum* | *Evylaeus* | 1 | | |
| Halictidae | *Lasioglossum* | *Lasioglossum* | *morrilli* | (Cockerell) | |
| Halictidae | *Lasioglossum* | *Lasioglossum* | *sisymbrii* | (Cockerell) | |
| Halictidae | *Lasioglossum* | *Sphecodogastra* | *lusorium* | (Cresson) | |
| Halictidae | *Nomia* | *Acunomia* | *tetrazonata uvaldensis* | Cockerell | |
| Halictidae | *Nomia* | *Epinomia* | *triangulifera* | Vachal | |
| Halictidae | *Sphecodes* | NA | 1 | | |

| Family | Genus | Subgenus | Species | Author | Notes |
|---|---|---|---|---|---|
| Halictidae | *Sphecodes* | NA | 2 | | |
| Halictidae | *Sphecodes* | NA | 4 | | |
| Halictidae | *Sphecodes* | NA | 5 | | |
| Halictidae | *Sphecodes* | NA | 6 | | |
| Halictidae | *Sphecodosoma* | NA | *dicksoni* | (Timberlake) | |
| Megachilidae | *Anthidium* | *Anthidium* | *cockerelli* | Schwarz | |
| Megachilidae | *Anthidium* | *Anthidium* | *emarginatum* | (Say) | |
| Megachilidae | *Anthidium* | *Anthidium* | *maculosum* | Cresson | |
| Megachilidae | *Anthidium* | *Anthidium* | *palmarum* | Cockerell | |
| Megachilidae | *Anthidium* | *Anthidium* | *porterae* | Cockerell | |
| Megachilidae | *Anthidium* | *Anthidium* | *schwarzi* | Gonzalez & Griswold | |
| Megachilidae | *Ashmeadiella* | *Arogochila* | *erema* | Michener | |
| Megachilidae | *Ashmeadiella* | *Ashmeadiella* | *bigeloveae* | (Cockerell) | |
| Megachilidae | *Ashmeadiella* | *Ashmeadiella* | *bucconis* | (Say) | |
| Megachilidae | *Ashmeadiella* | *Ashmeadiella* | *cactorum* | (Cockerell) | |
| Megachilidae | *Ashmeadiella* | *Ashmeadiella* | *difugita* | Michener | |
| Megachilidae | *Ashmeadiella* | *Ashmeadiella* | *gillettei* | Titus | |
| Megachilidae | *Ashmeadiella* | *Ashmeadiella* | *meliloti* | (Cockerell) | |
| Megachilidae | *Ashmeadiella* | *Ashmeadiella* | *occipitalis* | Michener | |
| Megachilidae | *Ashmeadiella* | *Ashmeadiella* | *opuntiae* | (Cockerell) | |
| Megachilidae | *Ashmeadiella* | *Ashmeadiella* | *vandykiella* | Michener | |
| Megachilidae | *Ashmeadiella* | *Isosmia* | *rubrella* | (Michener) | |
| Megachilidae | *Atoposmia* | *Eremosmia* | aff. *daleae* | | New species |
| Megachilidae | *Atoposmia* | *Eremosmia* | aff. *daleae* 2 | | New species |
| Megachilidae | *Coelioxys* | *Xerocoelioxys* | *hirsutissima* | Cockerell | |
| Megachilidae | *Coelioxys* | *Xerocoelioxys* | *mesae* | Cockerell | |
| Megachilidae | *Dianthidium* | *Dianthidium* | 1 | | New species |
| Megachilidae | *Dianthidium* | *Dianthidium* | *concinnum* | (Cresson) | |
| Megachilidae | *Dianthidium* | *Dianthidium* | *heterulkei* | Schwarz | |
| Megachilidae | *Dianthidium* | *Dianthidium* | *implicatum* | Timberlake | |
| Megachilidae | *Dianthidium* | *Dianthidium* | *parvum* | (Cresson) | |
| Megachilidae | *Dioxys* | NA | aff. *pomonae* | | |
| Megachilidae | *Dioxys* | NA | *pacificus* | Cockerell | |
| Megachilidae | *Dioxys* | NA | *productus* | (Cresson) | |
| Megachilidae | *Hoplitis* | *Alcidamea* | *biscutellae* | (Cockerell) | |
| Megachilidae | *Hoplitis* | *Alcidamea* | *grinnelli* | (Cockerell) | |
| Megachilidae | *Hoplitis* | *Proteriades* | aff. *torchioi* | | |
| Megachilidae | *Hoplitis* | *Proteriades* | *zuni* | (Parker) | |
| Megachilidae | *Lithurgopsis* | NA | *apicalis* | (Cresson) | |
| Megachilidae | *Megachile* | *Argyropile* | *parallela* | Smith | |
| Megachilidae | *Megachile* | *Argyropile* | *townsendiana* | Cockerell | |
| Megachilidae | *Megachile* | *Chelostomoides* | *lobatifrons* | Cockerell | |
| Megachilidae | *Megachile* | *Chelostomoides* | *prosopidis* | Cockerell | |
| Megachilidae | *Megachile* | *Chelostomoides* | *spinotulata* | Mitchell | |
| Megachilidae | *Megachile* | *Litomegachile* | *brevis* | Say | |
| Megachilidae | *Megachile* | *Litomegachile* | *lippiae* | Cockerell | |
| Megachilidae | *Megachile* | *Megachile* | *montivaga* | Cresson | |
| Megachilidae | *Megachile* | *Megachiloides* | aff. *dakotensis* | | |
| Megachilidae | *Megachile* | *Megachiloides* | *casadae* | Cockerell | |
| Megachilidae | *Megachile* | *Megachiloides* | *fucata* | Mitchell | |
| Megachilidae | *Megachile* | *Megachiloides* | *melanderi* | Mitchell | |
| Megachilidae | *Megachile* | *Megachiloides* | *mucorosa* | Cockerell | |
| Megachilidae | *Megachile* | *Megachiloides* | *sublaurita* | Mitchell | |
| Megachilidae | *Megachile* | *Megachiloides* | *umatillensis* | (Mitchell) | |
| Megachilidae | *Megachile* | *Megachiloides* | *xerophila* | Cockerell | |
| Megachilidae | *Megachile* | *Pseudocentron* | *sidalceae* | Cockerell | |
| Megachilidae | *Megachile* | *Sayapis* | *inimica inimica* | Cresson | |
| Megachilidae | *Megachile* | *Sayapis* | *policaris* | Say | |
| Megachilidae | *Osmia* | *Helicosmia* | *coloradensis* | Cresson | |
| Megachilidae | *Osmia* | *Melanosmia* | 1 | | |

| Family | Genus | Subgenus | Species | Author | Notes |
|---|---|---|---|---|---|
| Megachilidae | *Osmia* | *Melanosmia* | 2 | | |
| Megachilidae | *Osmia* | *Melanosmia* | aff. *enixa* | | New species |
| Megachilidae | *Osmia* | *Melanosmia* | *cerasi* | Cockerell | |
| Megachilidae | *Osmia* | *Melanosmia* | *cordata* | Robertson | |
| Megachilidae | *Osmia* | *Melanosmia* | *gaudiosa* | Cockerell | |
| Megachilidae | *Osmia* | *Melanosmia* | *integra* | Cresson | |
| Megachilidae | *Osmia* | *Melanosmia* | *liogastra* | Cockerell | |
| Megachilidae | *Osmia* | *Melanosmia* | *phenax* | Cockerell | |
| Megachilidae | *Osmia* | *Melanosmia* | *prunorum* | Cockerell | |
| Megachilidae | *Osmia* | *Melanosmia* | *trevoris* | Cockerell | |
| Megachilidae | *Osmia* | *Melanosmia* | *watsoni* | Cockerell | |
| Megachilidae | *Osmia* | *Trichinosmia* | *latisulcata* | Michener | |
| Megachilidae | *Stelis* | *Stelis* | *imperialis* | Parker & Griswold | |
| Megachilidae | *Stelis* | *Stelis* | *palmarum* | Timberlake | |
| Megachilidae | *Stelis* | *Stelis* | *robertsoni* | Timberlake | |
| Megachilidae | *Trachusa* | *Heteranthidium* | *larreae* | (Cockerell) | |
| Mellitidae | *Hesperapis* | *Amblyapis* | *larreae* | Cockerell | |
| Mellitidae | *Hesperapis* | *Hesperapis* | *trochanterata* | Snelling | |

**Supplementary Table S2.** Abundance and diversity measures for bees in three ecosystem types of the Sevilleta National Wildlife Refuge, collected in funnel traps, 2002–2014. "Overall" measures were calculated from data pooled across years, months, and transects within an ecosystem.

| | Plains grassland | Chihuahuan Desert grassland | Chihuahuan Desert shrubland |
|---|---|---|---|
| Overall abundance | 26,462 | 26,872 | 17,617 |
| Mean monthly per-transect abundance | 50.89 | 51.68 | 33.88 |
| Overall richness | 217 | 207 | 216 |
| Mean monthly per-transect richness | 32.90 | 31.13 | 35.50 |
| Overall Shannon $H'$ | 2.58 | 2.11 | 2.69 |
| Mean monthly per-transect Shannon $H'$ | 1.99 | 1.71 | 2.08 |
| Overall Pielou's $J$ | 0.479 | 0.395 | 0.500 |
| Mean monthly per-transect Pielou's $J$ | 0.578 | 0.516 | 0.592 |

**Supplementary Table S3.** Indicator species of each month (March–October) within each of three ecosystem types (Plains grassland, Chihuahuan Desert grassland, and Chihuahuan Desert shrubland), according to Dufrene-Legendre indicator species value. Grey highlights indicate cases in which two or more ecosystem types shared an indicator species in a given month. Asterisks (*) signify cases in which bee species were indicators of different months in different ecosystem types.

| | Plains grassland | Chihuahuan Desert grassland | Chihuahuan Desert shrubland |
|---|---|---|---|
| March | *Andrena prima* | *Andrena prima* | *Andrena prima* |
| | *Anthophora porterae* | *Anthophora porterae* | *Anthophora porterae* |
| | *Dioxys pacificus* | *Dioxys pacificus* | *Dioxys pacificus* |
| | *Eucera lycii* | *Eucera lycii* | *Eucera lycii* |
| | *Eucera* aff. *albescens* | *Eucera* aff. *albescens* | *Eucera* aff. *albescens* |
| | *Halictus tripartitus* | *Halictus tripartitus* | *Halictus tripartitus* |
| | *Melecta pacifica* | *Melecta pacifica* | *Melecta pacifica* |
| | *Osmia prunorum* | *Osmia prunorum* | *Osmia prunorum* |
| | *Osmia watsoni* | *Osmia watsoni* | *Osmia watsoni* |
| | | *Anthidium emarginatum** | *Anthidium emarginatum** |
| | | *Anthophora lesquerellae** | *Anthophora lesquerellae** |
| | | *Ashmeadiella erema* | *Ashmeadiella erema* |
| | | *Osmia phenax* | *Osmia phenax* |
| | *Eucera territella* | *Eucera territella* | |
| | *Habropoda morrisoni* | *Habropoda morrisoni* | |
| | *Osmia* aff. *enixa* | *Osmia* aff. *enixa* | |
| | *Osmia cerasi** | *Osmia cerasi** | |
| | *Osmia liogastra* | *Osmia liogastra* | |
| | *Dioxys* aff. *pomonae* | | *Dioxys* aff. *pomonae* |
| | *Lasioglossum sisymbrii** | *Andrena mesillae* | *Andrena illinoiensis* |
| | *Osmia* (*Melanosmia*) sp. 2 | *Atoposmia* aff. *daleae* 2 | *Anthidium cockerelli** |
| | *Osmia cordata* | | *Ashmeadiella rubrella* |
| | *Osmia latisulcata* | | *Lasioglossum semicaeruleum** |
| | | | *Melecta alexanderi** |
| | | | *Melecta bohartorum** |
| April | *Anthophora affabilis* | *Anthophora affabilis* | *Anthophora affabilis* |
| | *Anthophora californica* | *Anthophora californica* | *Anthophora californica* |
| | *Anthophora* n. sp. | *Anthophora* n. sp. | *Anthophora* n. sp. |
| | *Lasioglossum* (*Dialictus*) sp. 2 | *Lasioglossum* (*Dialictus*) sp. 2 | *Lasioglossum* (*Dialictus*) sp. 2 |
| | | *Andrena* (*Micrandrena*) sp. 2 | *Andrena* (*Micrandrena*) sp. 2 |
| | | *Lasioglossum sisymbrii** | *Lasioglossum sisymbrii** |
| | | *Megachile sublaurita* | *Megachile sublaurita* |
| | *Anthidium schwarzi* | *Anthidium schwarzi* | |
| | *Dufourea pulchricornis* | *Dufourea pulchricornis* | |
| | *Melecta alexanderi** | *Melecta alexanderi** | |

| | Plains grassland | Chihuahuan Desert grassland | Chihuahuan Desert shrubland |
|---|---|---|---|
| | *Melecta bohartorum** | *Melecta bohartorum** | |
| | *Agapostemon angelicus** | *Anthidium cockerelli** | *Ashmeadiella bucconis* |
| | *Anthidium emarginatum** | *Atoposmia* aff. *daleae* | *Ashmeadiella cactorum* |
| | *Anthophora lesquerellae** | *Hylaeus episcopelis coquilletti* | *Dufourea vernalis* |
| | *Coelioxys hirsutissima* | *Lasioglossum semicaeruleum** | *Osmia cerasi** |
| | *Dioxys productus* | *Megachile fucata* | |
| | *Lasioglossum hudsoniellum** | | |
| May | *Perdita* (*Perdita*) sp. 36 | *Perdita* (*Perdita*) sp. 36 | *Perdita* (*Perdita*)sp. 36 |
| | | *Hoplitis biscutellae* | *Hoplitis biscutellae* |
| | | *Perdita coreopsidis kansensis* | *Perdita coreopsidis kansensis* |
| | *Lasioglossum morrilli* | | *Lasioglossum morrilli* |
| | *Ceratina nanula* | *Anthophora montana** | |
| | *Megachile policaris* | *Colletes clypeonitens* | |
| | | *Lithurgus apicalis** | |
| June | *Diadasia australis* | *Diadasia australis* | *Diadasia australis* |
| | *Diadasia rinconis* | *Diadasia rinconis* | *Diadasia rinconis* |
| | *Anthophora urbana* | *Agapostemon angelicus** | *Lasioglossum* (*Dialictus*) sp. 5 |
| | *Colletes scopiventer* | *Lasioglossum* (*Dialictus*) sp. 7* | *Lasioglossum* (*Dialictus*) sp. 8 |
| | *Lithurgus apicalis** | | *Lasioglossum* (*Dialictus*) sp. 13 |
| | *Melissodes paroselae* | | *Lasioglossum hudsoniellum** |
| | | | *Lasioglossum microlepoides** |
| | | | *Perdita fallax** |
| July | *Halictus ligatus* | *Halictus ligatus* | *Halictus ligatus* |
| | *Martinapis luteicornis* | *Martinapis luteicornis* | *Martinapis luteicornis* |
| | *Melissodes tristis* | *Melissodes tristis* | *Melissodes tristis* |
| | | *Ashmeadiella meliloti* | *Ashmeadiella meliloti* |
| | | *Diadasia ochracea** | *Diadasia ochracea** |
| | | *Lasioglossum comulum* | *Lasioglossum comulum* |
| | | *Perdita callicerata** | *Perdita callicerata** |
| | | *Perdita diversa* | *Perdita diversa* |
| | | *Perdita ignota ignota** | *Perdita ignota ignota** |
| | *Agapostemon melliventris** | *Agapostemon melliventris** | |
| | *Lasioglossum* aff. *pervarum* | *Lasioglossum* aff. *pervarum* | |
| | *Anthidium porterae** | *Anthophorula compactula** | *Perdita albovittata* |
| | *Lasioglossum* (*Dialictus*) sp. 7* | *Lasioglossum* (*Dialictus*) sp. 11 | |
| | *Lasioglossum microlepoides** | *Lasioglossum lusoria* | |
| | *Lasioglossum semicaeruleum** | *Neolarra vigilans** | |
| | *Melissodes thelypodii thelypodii* | *Perdita fallax** | |

| | Plains grassland | Chihuahuan Desert grassland | Chihuahuan Desert shrubland |
|---|---|---|---|
| | *Sphecodes* sp. 5 | | |
| August | *Perdita larreae* | *Perdita larreae* | *Perdita larreae* |
| | *Perdita marcialis* | *Perdita marcialis* | *Perdita marcialis* |
| | *Perdita semicaerulea* | *Perdita semicaerulea* | *Perdita semicaerulea* |
| | | *Hesperapis larreae* | *Hesperapis larreae* |
| | | *Perdita drymariae* | *Perdita drymariae* |
| | *Anthophora montana** | *Anthidium porterae** | *Agapostemon melliventris** |
| | *Diadasia ochracea** | *Melissodes communis communis* | *Anthophorula compactula** |
| | *Melissodes coreopsis* | *Svastra helianthelli* | *Macrotera portalis** |
| | *Perdita callicerata** | | *Melissodes agilis** |
| | *Perdita ignota ignota** | | *Neolarra vigilans** |
| | *Sphecodes* sp. 1 | | *Perdita aperta* |
| September | | *Diadasia diminuta* | *Diadasia diminuta* |
| | | *Macrotera latior* | *Macrotera latior* |
| | *Melissodes agilis** | *Melissodes agilis** | |
| | *Perdita* sp. 13 | *Anthophorula pygmaea* | *Anthophora montana** |
| | *Protandrena* sp. 2 | *Macrotera portalis** | *Anthophorula completa* |
| | | | *Diadasia megamorpha** |
| | | | *Macrotera magniceps* |
| October | None | *Diadasia megamorpha** | *Melissodes snowii* |
| | | | *Perdita austini* |

**Supplementary Table S4.** Indicator species for each month (March-October) within the Plains grassland ecosystem, according to Dufrene-Legendre indicator species value. Species are listed from highest to lowest indicator value within each month.

Plains grassland

| Species | Family | Indicator value | *P*-value |
|---|---|---|---|
| March | | | |
| *Osmia watsoni* | Megachilidae | 1.00 | 0.001 |
| *Eucera lycii* | Apidae | 0.99 | 0.001 |
| *Anthophora porterae* | Apidae | 0.99 | 0.001 |
| *Melecta pacifica* | Apidae | 0.98 | 0.001 |
| *Osmia* aff. *enixa* | Megachilidae | 0.95 | 0.001 |
| *Eucera* aff. *albescens* | Apidae | 0.83 | 0.001 |
| *Osmia cerasi* | Megachilidae | 0.82 | 0.001 |
| *Osmia cordata* | Megachilidae | 0.80 | 0.003 |
| *Osmia latisulcata* | Megachilidae | 0.80 | 0.001 |
| *Osmia liogastra* | Megachilidae | 0.80 | 0.001 |
| *Dioxys pacificus* | Megachilidae | 0.74 | 0.001 |
| *Andrena prima* | Andrenidae | 0.60 | 0.011 |
| *Eucera territella* | Apidae | 0.58 | 0.002 |
| *Osmia* (*Melanosmia*) sp. 2 | Megachilidae | 0.54 | 0.009 |
| *Osmia prunorum* | Megachilidae | 0.50 | 0.004 |
| *Dioxys* aff. *pomonae* | Megachilidae | 0.49 | 0.010 |
| *Habropoda morrisoni* | Apidae | 0.45 | 0.016 |
| *Halictus tripartitus* | Halictidae | 0.40 | 0.008 |
| *Lasioglossum sisymbrii* | Halictidae | 0.34 | 0.002 |
| April | | | |
| *Dufourea pulchricornis* | Halictidae | 1.00 | 0.001 |
| *Coelioxys hirsutissima* | Megachilidae | 1.00 | 0.001 |
| *Anthophora affabilis* | Apidae | 0.84 | 0.001 |
| *Melecta alexanderi* | Apidae | 0.78 | 0.001 |
| *Lasioglossum* (*Dialictus*) sp. 2 | Halictidae | 0.77 | 0.001 |
| *Anthidium emarginatum* | Megachilidae | 0.65 | 0.001 |
| *Melecta bohartorum* | Apidae | 0.63 | 0.002 |
| *Anthophora* n. sp. | Apidae | 0.60 | 0.013 |
| *Dioxys productus* | Megachilidae | 0.60 | 0.011 |
| *Anthophora lesquerellae* | Apidae | 0.53 | 0.001 |
| *Anthidium schwarzi* | Megachilidae | 0.52 | 0.002 |
| *Lasioglossum hudsoniellum* | Halictidae | 0.42 | 0.001 |
| *Anthophora californica* | Apidae | 0.41 | 0.014 |
| *Agapostemon angelicus* | Halictidae | 0.22 | 0.001 |
| May | | | |
| *Lasioglossum morrilli* | Halictidae | 0.62 | 0.001 |
| *Perdita* (*Perdita*) sp. 36 | Andrenidae | 0.60 | 0.006 |
| *Ceratina nanula* | Apidae | 0.60 | 0.009 |
| *Megachile policaris* | Megachilidae | 0.36 | 0.050 |
| June | | | |
| *Diadasia australis* | Apidae | 0.65 | 0.001 |
| *Lithurgus apicalis* | Megachilidae | 0.57 | 0.003 |
| *Colletes scopiventer* | Colletidae | 0.46 | 0.018 |
| *Diadasia rinconis* | Apidae | 0.42 | 0.001 |
| *Melissodes paroselae* | Apidae | 0.36 | 0.038 |
| *Anthophora urbana* | Apidae | 0.34 | 0.033 |
| July | | | |
| *Melissodes thelypodii thelypodii* | Apidae | 0.80 | 0.001 |

| | | | |
|---|---|---|---|
| *Martinapis luteicornis* | Apidae | 0.71 | 0.002 |
| *Anthidium porterae* | Megachilidae | 0.61 | 0.001 |
| *Agapostemon melliventris* | Halictidae | 0.56 | 0.004 |
| *Lasioglossum microlepoides* | Halictidae | 0.47 | 0.005 |
| *Lasioglossum* (*Dialictus*) sp. 7 | Halictidae | 0.37 | 0.040 |
| *Sphecodes* sp. 5 | Halictidae | 0.36 | 0.046 |
| *Halictus ligatus* | Halictidae | 0.33 | 0.002 |
| *Lasioglossum* aff. *pervarum* | Halictidae | 0.32 | 0.002 |
| *Melissodes tristis* | Apidae | 0.32 | 0.001 |
| *Lasioglossum semicaeruleum* | Halictidae | 0.20 | 0.023 |
| August | | | |
| *Perdita semicaerulea* | Andrenidae | 0.98 | 0.001 |
| *Perdita marcialis* | Andrenidae | 0.60 | 0.013 |
| *Melissodes coreopsis* | Apidae | 0.60 | 0.012 |
| *Perdita ignota ignota* | Andrenidae | 0.60 | 0.001 |
| *Perdita larreae* | Andrenidae | 0.51 | 0.012 |
| *Sphecodes* sp. 1 | Halictidae | 0.45 | 0.006 |
| *Anthophora montana* | Apidae | 0.40 | 0.001 |
| *Diadasia ochracea* | Apidae | 0.38 | 0.018 |
| *Perdita callicerata* | Andrenidae | 0.36 | 0.004 |
| September | | | |
| *Perdita* sp. 13 | Andrenidae | 0.60 | 0.013 |
| *Melissodes agilis* | Apidae | 0.51 | 0.003 |
| *Protandrena* sp. 2 | Andrenidae | 0.45 | 0.030 |
| October | | | |
| None | | | |

**Supplementary Table S5.** Indicator species for each month (March-October) within the Chihuahuan Desert grassland ecosystem, according to Dufrene-Legendre indicator species value. Species are listed from highest to lowest indicator value within each month.

Chihuahuan Desert grassland

| Species | Family | Indicator value | *P*-value |
|---|---|---|---|
| March | | | |
| *Eucera lycii* | Apidae | 0.98 | 0.001 |
| *Osmia watsoni* | Megachilidae | 0.98 | 0.001 |
| *Anthophora porterae* | Apidae | 0.96 | 0.001 |
| *Halictus tripartitus* | Halictidae | 0.96 | 0.001 |
| *Melecta pacifica* | Apidae | 0.95 | 0.001 |
| *Andrena prima* | Andrenidae | 0.88 | 0.001 |
| *Osmia phenax* | Megachilidae | 0.84 | 0.002 |
| *Eucera* aff. *albescens* | Apidae | 0.80 | 0.002 |
| *Anthidium emarginatum* | Megachilidae | 0.75 | 0.001 |
| *Osmia* aff. *enixa* | Megachilidae | 0.73 | 0.001 |
| *Osmia liogastra* | Megachilidae | 0.70 | 0.001 |
| *Andrena mesillae* | Andrenidae | 0.60 | 0.009 |
| *Dioxys pacificus* | Megachilidae | 0.60 | 0.004 |
| *Osmia prunorum* | Megachilidae | 0.59 | 0.003 |
| *Anthophora lesquerellae* | Apidae | 0.58 | 0.001 |
| *Osmia cerasi* | Megachilidae | 0.57 | 0.004 |
| *Ashmeadiella erema* | Megachilidae | 0.48 | 0.036 |
| *Habropoda morrisoni* | Apidae | 0.47 | 0.007 |
| *Atoposmia* aff. *daleae* 2 | Megachilidae | 0.45 | 0.031 |
| *Eucera territella* | Apidae | 0.42 | 0.015 |
| April | | | |
| *Anthophora* n. sp. | Apidae | 0.90 | 0.001 |
| *Anthophora californica* | Apidae | 0.80 | 0.001 |
| *Melecta alexanderi* | Apidae | 0.76 | 0.001 |
| *Anthophora affabilis* | Apidae | 0.76 | 0.001 |
| *Andrena* (*Micrandrena*) sp. 2 | Andrenidae | 0.74 | 0.001 |
| *Megachile sublaurita* | Megachilidae | 0.66 | 0.001 |
| *Melecta bohartorum* | Apidae | 0.63 | 0.001 |
| *Hylaeus episcopelis coquilletti* | Colletidae | 0.60 | 0.012 |
| *Anthidium cockerelli* | Megachilidae | 0.60 | 0.013 |
| *Megachile fucata* | Megachilidae | 0.60 | 0.007 |
| *Lasioglossum sisymbrii* | Halictidae | 0.58 | 0.001 |
| *Dufourea pulchricornis* | Halictidae | 0.56 | 0.009 |
| *Lasioglossum* (*Dialictus*) sp. 2 | Halictidae | 0.55 | 0.001 |
| *Anthidium schwarzi* | Megachilidae | 0.53 | 0.013 |
| *Atoposmia* aff. *daleae* | Megachilidae | 0.45 | 0.029 |
| *Lasioglossum semicaeruleum* | Halictidae | 0.24 | 0.003 |
| May | | | |
| *Hoplitis biscutellae* | Megachilidae | 0.80 | 0.001 |
| *Perdita* (*Perdita*) sp. 36 | Andrenidae | 0.73 | 0.003 |
| *Colletes clypeonitens* | Colletidae | 0.60 | 0.011 |
| *Lithurgus apicalis* | Megachilidae | 0.51 | 0.015 |
| *Perdita coreopsidis kansensis* | Andrenidae | 0.45 | 0.001 |
| *Anthophora montana* | Apidae | 0.41 | 0.013 |
| June | | | |
| *Diadasia australis* | Apidae | 0.84 | 0.001 |
| *Diadasia rinconis* | Apidae | 0.46 | 0.002 |
| *Lasioglossum* (*Dialictus*) sp. 7 | Halictidae | 0.44 | 0.007 |
| *Agapostemon angelicus* | Halictidae | 0.23 | 0.009 |

| | | | |
|---|---|---|---|
| July | | | |
| *Lasioglossum* (*Dialictus*) sp. 11 | Halictidae | 0.89 | 0.001 |
| *Lasioglossum comulum* | Halictidae | 0.74 | 0.001 |
| *Diadasia ochracea* | Apidae | 0.70 | 0.002 |
| *Perdita ignota ignota* | Andrenidae | 0.69 | 0.001 |
| *Martinapis luteicornis* | Apidae | 0.58 | 0.005 |
| *Agapostemon melliventris* | Halictidae | 0.58 | 0.001 |
| *Halictus ligatus* | Halictidae | 0.57 | 0.001 |
| *Perdita callicerata* | Andrenidae | 0.55 | 0.001 |
| *Neolarra vigilans* | Apidae | 0.48 | 0.020 |
| *Lasioglossum* aff. *pervarum* | Halictidae | 0.45 | 0.001 |
| *Lasioglossum lusoria* | Halictidae | 0.42 | 0.011 |
| *Ashmeadiella meliloti* | Megachilidae | 0.37 | 0.001 |
| *Anthophorula compactula* | Apidae | 0.37 | 0.027 |
| *Perdita diversa* | Andrenidae | 0.37 | 0.029 |
| *Melissodes tristis* | Apidae | 0.34 | 0.001 |
| *Perdita fallax* | Andrenidae | 0.30 | 0.024 |
| August | | | |
| *Hesperapis larreae* | Mellitidae | 1.00 | 0.001 |
| *Perdita semicaerulea* | Andrenidae | 0.93 | 0.001 |
| *Perdita drymariae* | Andrenidae | 0.89 | 0.001 |
| *Perdita marcialis* | Andrenidae | 0.85 | 0.001 |
| *Perdita larreae* | Andrenidae | 0.85 | 0.001 |
| *Melissodes communis communis* | Apidae | 0.71 | 0.001 |
| *Svastra helianthelli* | Apidae | 0.60 | 0.011 |
| *Anthidium porterae* | Megachilidae | 0.58 | 0.001 |
| September | | | |
| *Macrotera portalis* | Andrenidae | 0.87 | 0.001 |
| *Anthophorula pygmaea* | Apidae | 0.80 | 0.002 |
| *Melissodes agilis* | Apidae | 0.48 | 0.007 |
| *Diadasia diminuta* | Apidae | 0.40 | 0.003 |
| *Macrotera latior* | Andrenidae | 0.38 | 0.034 |
| October | | | |
| *Diadasia megamorpha* | Apidae | 0.53 | 0.003 |

**Supplementary Table S6**. Indicator species for each month (March-October) within the Chihuahuan Desert shrubland ecosystem, according to Dufrene-Legendre indicator species value. Species are listed from highest to lowest indicator value within each month.

Chihuahuan Desert shrubland

| Species | Family | Indicator value | *P*-value |
|---|---|---|---|
| March | | | |
| *Osmia phenax* | Megachilidae | 1.00 | 0.001 |
| *Melecta pacifica* | Apidae | 1.00 | 0.001 |
| *Eucera lycii* | Apidae | 0.99 | 0.001 |
| *Anthophora porterae* | Apidae | 0.98 | 0.001 |
| *Osmia watsoni* | Megachilidae | 0.96 | 0.001 |
| *Andrena illinoiensis* | Andrenidae | 0.80 | 0.002 |
| *Melecta alexanderi* | Apidae | 0.80 | 0.002 |
| *Dioxys* aff. *pomonae* | Megachilidae | 0.78 | 0.002 |
| *Ashmeadiella erema* | Megachilidae | 0.74 | 0.002 |
| *Eucera* aff. *albescens* | Apidae | 0.72 | 0.003 |
| *Anthidium emarginatum* | Megachilidae | 0.70 | 0.001 |
| *Halictus tripartitus* | Halictidae | 0.69 | 0.001 |
| *Andrena prima* | Andrenidae | 0.60 | 0.006 |
| *Ashmeadiella rubrella* | Megachilidae | 0.60 | 0.006 |
| *Dioxys pacificus* | Megachilidae | 0.60 | 0.006 |
| *Anthophora lesquerellae* | Apidae | 0.57 | 0.001 |
| *Melecta bohartorum* | Apidae | 0.53 | 0.003 |
| *Osmia prunorum* | Megachilidae | 0.51 | 0.010 |
| *Anthidium cockerelli* | Megachilidae | 0.50 | 0.004 |
| *Lasioglossum semicaeruleum* | Halictidae | 0.29 | 0.001 |
| April | | | |
| *Anthophora affabilis* | Apidae | 0.84 | 0.001 |
| *Megachile sublaurita* | Megachilidae | 0.67 | 0.001 |
| *Dufourea vernalis* | Halictidae | 0.62 | 0.003 |
| *Ashmeadiella bucconis* | Megachilidae | 0.60 | 0.013 |
| *Ashmeadiella cactorum* | Megachilidae | 0.55 | 0.002 |
| *Andrena* (*Micrandrena*) sp. 2 | Andrenidae | 0.55 | 0.004 |
| *Anthophora* n. sp. | Apidae | 0.54 | 0.007 |
| *Osmia cerasi* | Megachilidae | 0.48 | 0.010 |
| *Anthophora californica* | Apidae | 0.42 | 0.009 |
| *Lasioglossum sisymbrii* | Halictidae | 0.39 | 0.004 |
| *Lasioglossum* (*Dialictus*) sp. 2 | Halictidae | 0.36 | 0.031 |
| May | | | |
| *Hoplitis biscutellae* | Megachilidae | 0.58 | 0.005 |
| *Lasioglossum morrilli* | Halictidae | 0.48 | 0.001 |
| *Perdita coreopsidis kansensis* | Andrenidae | 0.41 | 0.006 |
| *Perdita* (*Perdita*) sp. 36 | Andrenidae | 0.36 | 0.030 |
| June | | | |
| *Diadasia australis* | Apidae | 0.83 | 0.001 |
| *Lasioglossum* (*Dialictus*) sp. 5 | Halictidae | 0.60 | 0.008 |
| *Lasioglossum* (*Dialictus*) sp. 13 | Halictidae | 0.60 | 0.005 |
| *Lasioglossum microlepoides* | Halictidae | 0.47 | 0.001 |
| *Diadasia rinconis* | Apidae | 0.44 | 0.001 |
| *Lasioglossum* (*Dialictus*) sp. 8 | Halictidae | 0.38 | 0.033 |
| *Perdita fallax* | Andrenidae | 0.31 | 0.027 |
| *Lasioglossum hudsoniellum* | Halictidae | 0.31 | 0.002 |
| July | | | |
| *Martinapis luteicornis* | Apidae | 0.77 | 0.001 |

| | | | |
|---|---|---|---|
| *Diadasia ochracea* | Apidae | 0.73 | 0.001 |
| *Lasioglossum comulum* | Halictidae | 0.63 | 0.004 |
| *Perdita ignota ignota* | Andrenidae | 0.44 | 0.006 |
| *Perdita callicerata* | Andrenidae | 0.43 | 0.001 |
| *Perdita diversa* | Andrenidae | 0.42 | 0.011 |
| *Halictus ligatus* | Halictidae | 0.41 | 0.004 |
| *Perdita albovittata* | Andrenidae | 0.40 | 0.036 |
| *Ashmeadiella meliloti* | Megachilidae | 0.30 | 0.009 |
| *Melissodes tristis* | Apidae | 0.25 | 0.001 |
| August | | | |
| *Hesperapis larreae* | Mellitidae | 0.99 | 0.001 |
| *Perdita larreae* | Andrenidae | 0.94 | 0.001 |
| *Perdita semicaerulea* | Andrenidae | 0.88 | 0.001 |
| *Perdita marcialis* | Andrenidae | 0.81 | 0.001 |
| *Perdita drymariae* | Andrenidae | 0.65 | 0.003 |
| *Macrotera portalis* | Andrenidae | 0.61 | 0.002 |
| *Perdita aperta* | Andrenidae | 0.60 | 0.009 |
| *Anthophorula compactula* | Apidae | 0.57 | 0.001 |
| *Neolarra vigilans* | Apidae | 0.46 | 0.009 |
| *Melissodes agilis* | Apidae | 0.38 | 0.026 |
| *Agapostemon melliventris* | Halictidae | 0.33 | 0.027 |
| September | | | |
| *Macrotera latior* | Andrenidae | 0.45 | 0.010 |
| *Macrotera magniceps* | Andrenidae | 0.45 | 0.039 |
| *Diadasia diminuta* | Apidae | 0.41 | 0.001 |
| *Anthophora montana* | Apidae | 0.39 | 0.002 |
| *Diadasia megamorpha* | Apidae | 0.36 | 0.038 |
| *Anthophorula completa* | Apidae | 0.33 | 0.041 |
| October | | | |
| *Melissodes snowii* | Apidae | 0.40 | 0.023 |
| *Perdita austini* | Andrenidae | 0.34 | 0.042 |

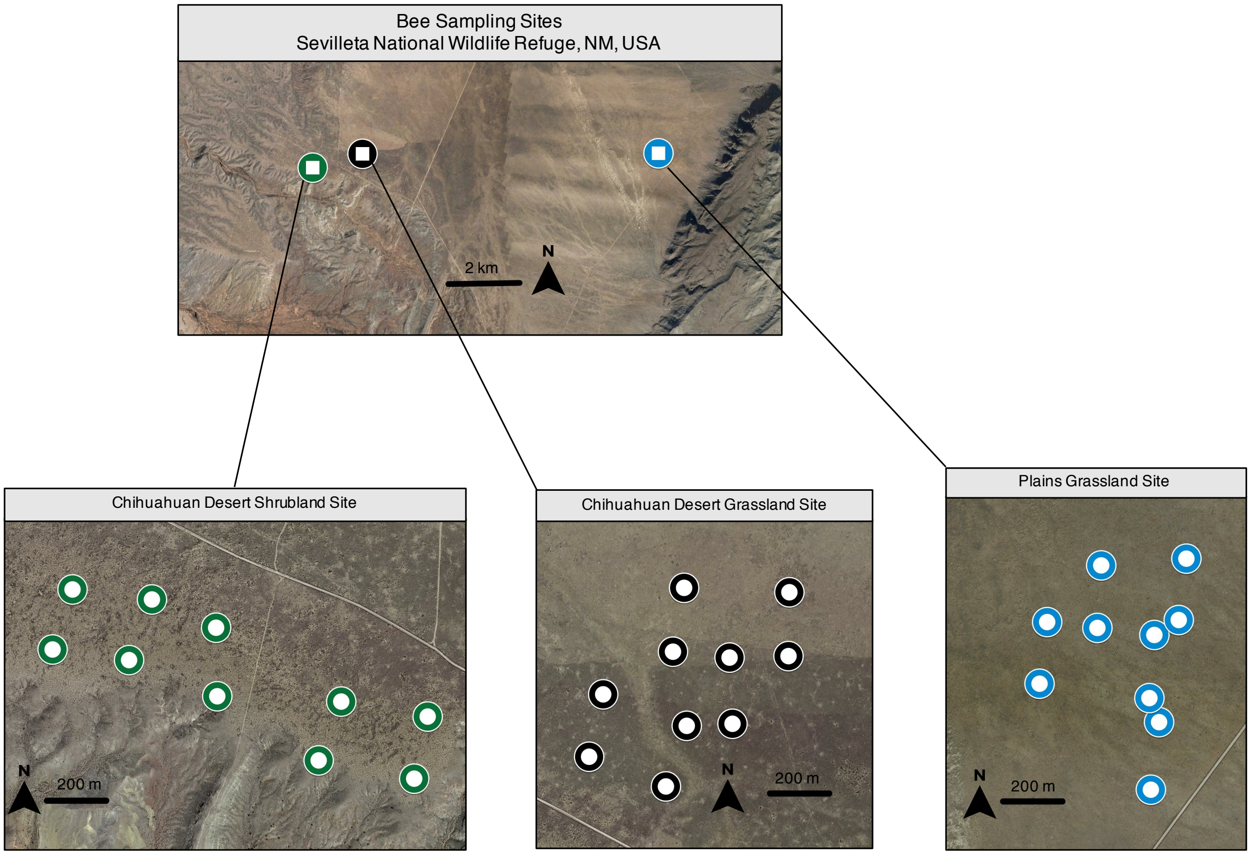


**Supplementary Figure S1.** Map of sampling sites at the Sevilleta National Wildlife Refuge, NM, USA. Bees were sampled in three focal ecosystem types: Chihuahuan Desert shrubland, Chihuahuan Desert grassland, and Plains grassland. To sample bees, we installed one passive funnel trap at each end of five 200 m transects/site; traps are indicated by colored points on the three lower panels.

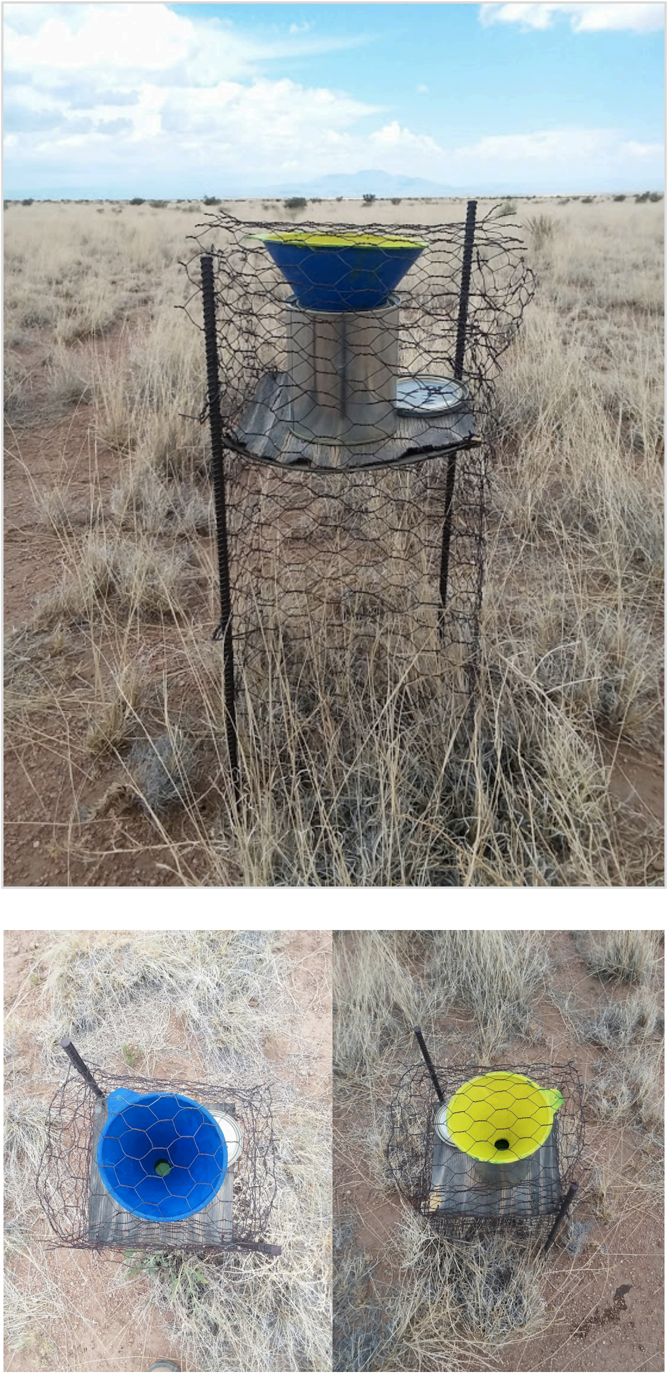

**Supplementary Figure S2.** Funnel traps used for bee collection. Each trap consisted of a 946 mL paint can filled with ~275 mL of propylene glycol and topped with a plastic automotive funnel (funnel height = 10 cm, top diameter = 14 cm, bottom diameter = 2.5 cm). The funnels' interiors were painted with either blue or yellow fluorescent paint (Krylon, Cleveland, OH or Ace Hardware, Oak Brook, IL). Each trap was placed on a 45 cm high platform that was surrounded by a 60 cm high chicken wire cage to prevent wildlife and wind disturbance.

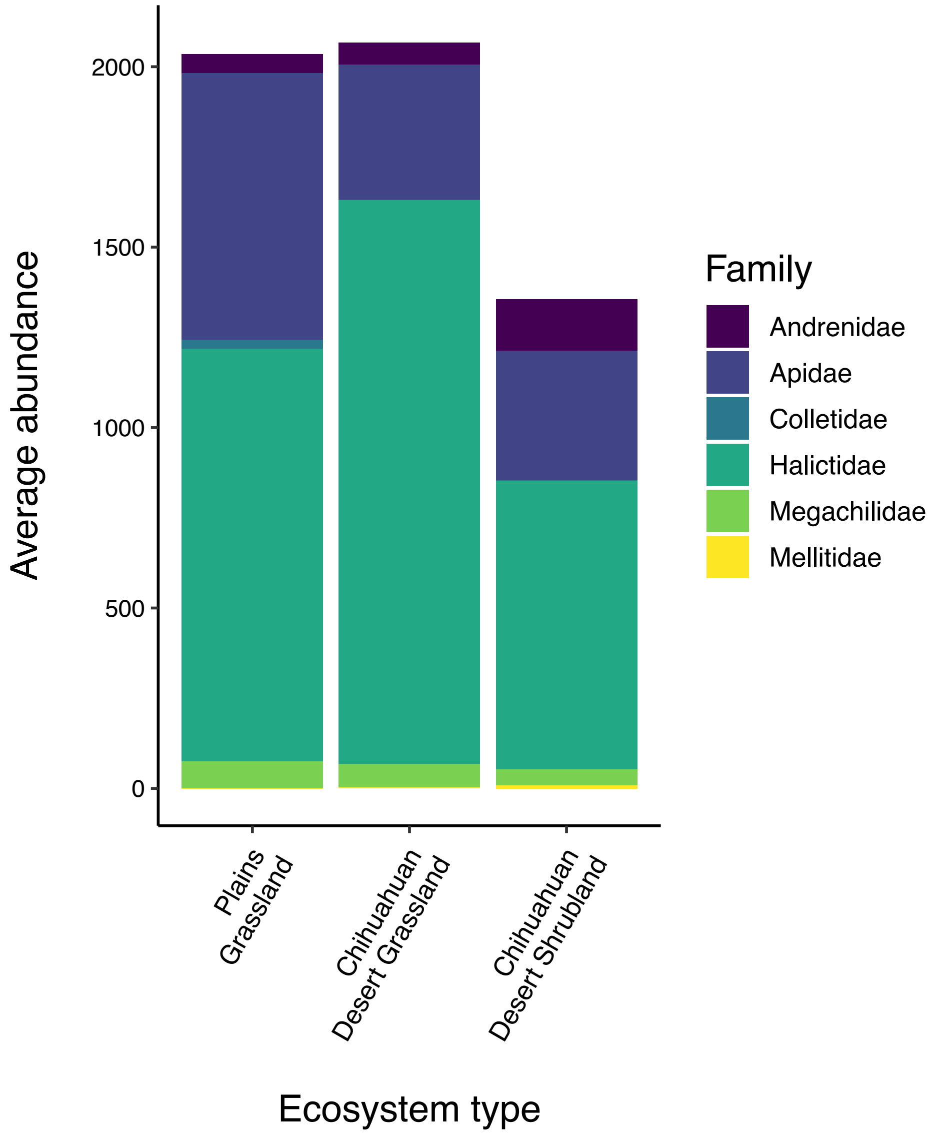


**Supplementary Figure S3**. Average yearly abundance of bees by family in each of the three studied southwestern U.S. ecosystem types.

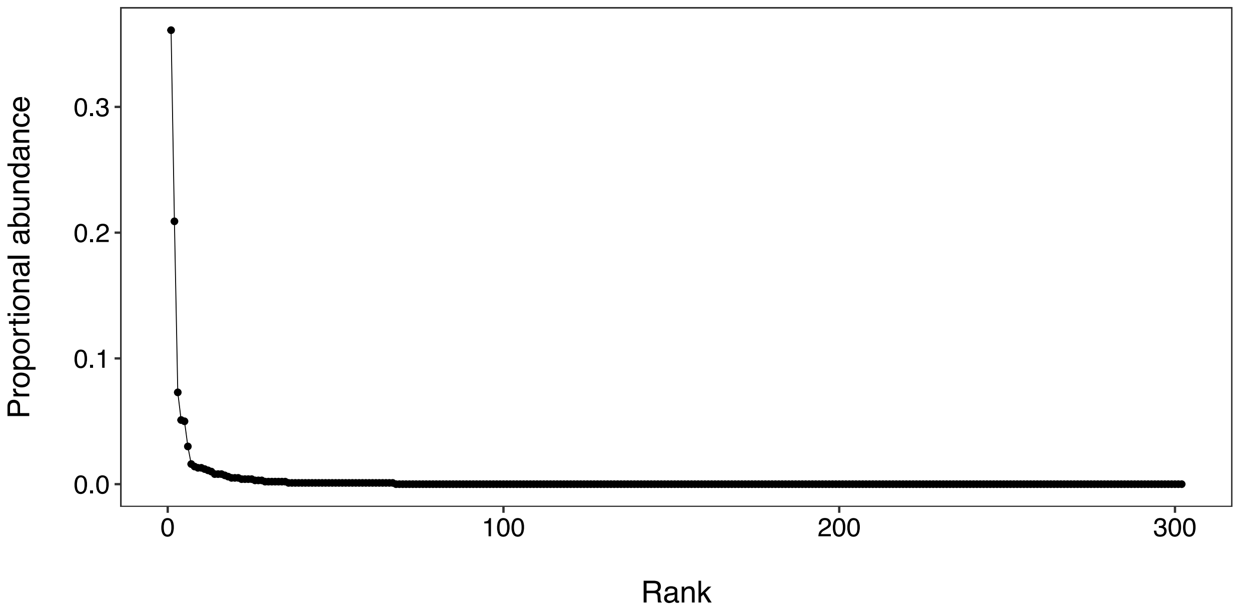


**Supplementary Figure S4.** Proportional abundance by abundance rank for the 302 bee species collected across three southwestern U.S. ecosystem types during the study.

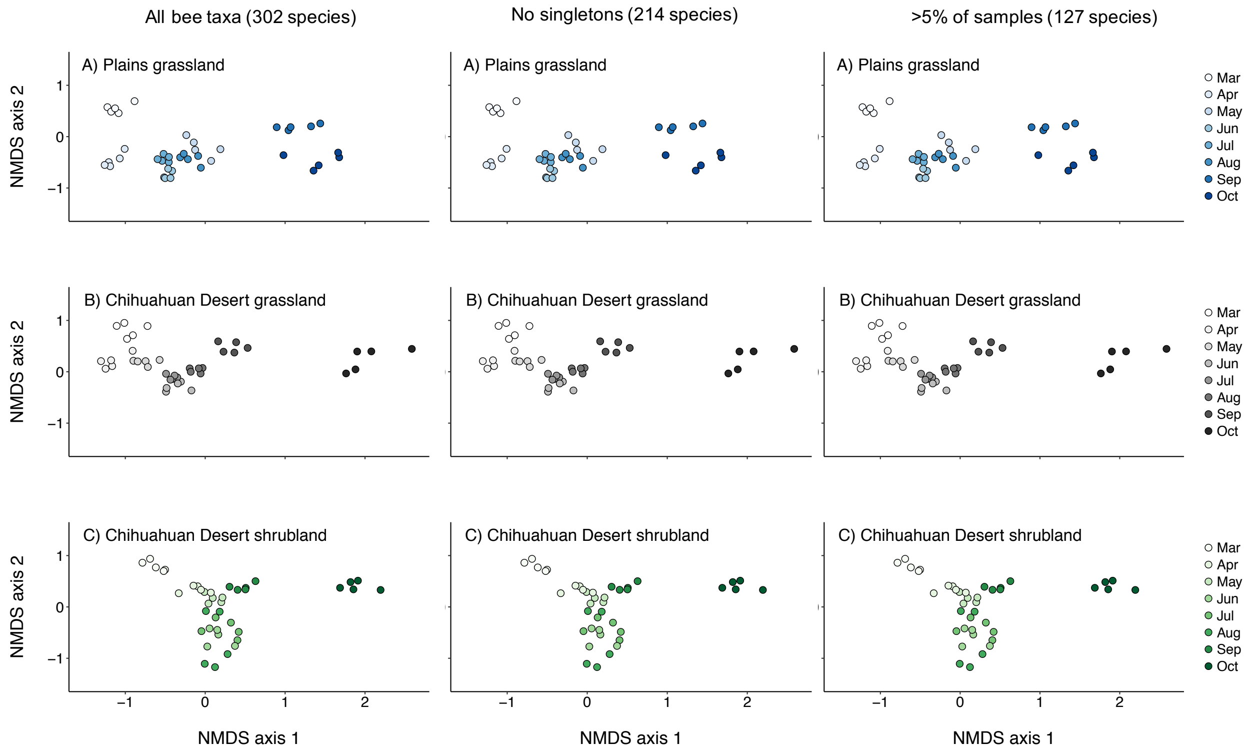


**Supplementary Figure S5.** Non-metric multidimensional scaling (NMDS) plots depicting variation in bee species composition for three dryland ecosystem types: (A) Plains grassland, (B) Chihuahuan Desert grassland, and (C) Chihuahuan Desert shrubland. NMDS analyses were run separately for the full dataset (left column), the dataset with singleton bee species removed (center column), and the dataset with only bee species present in >5% of samples (right column).

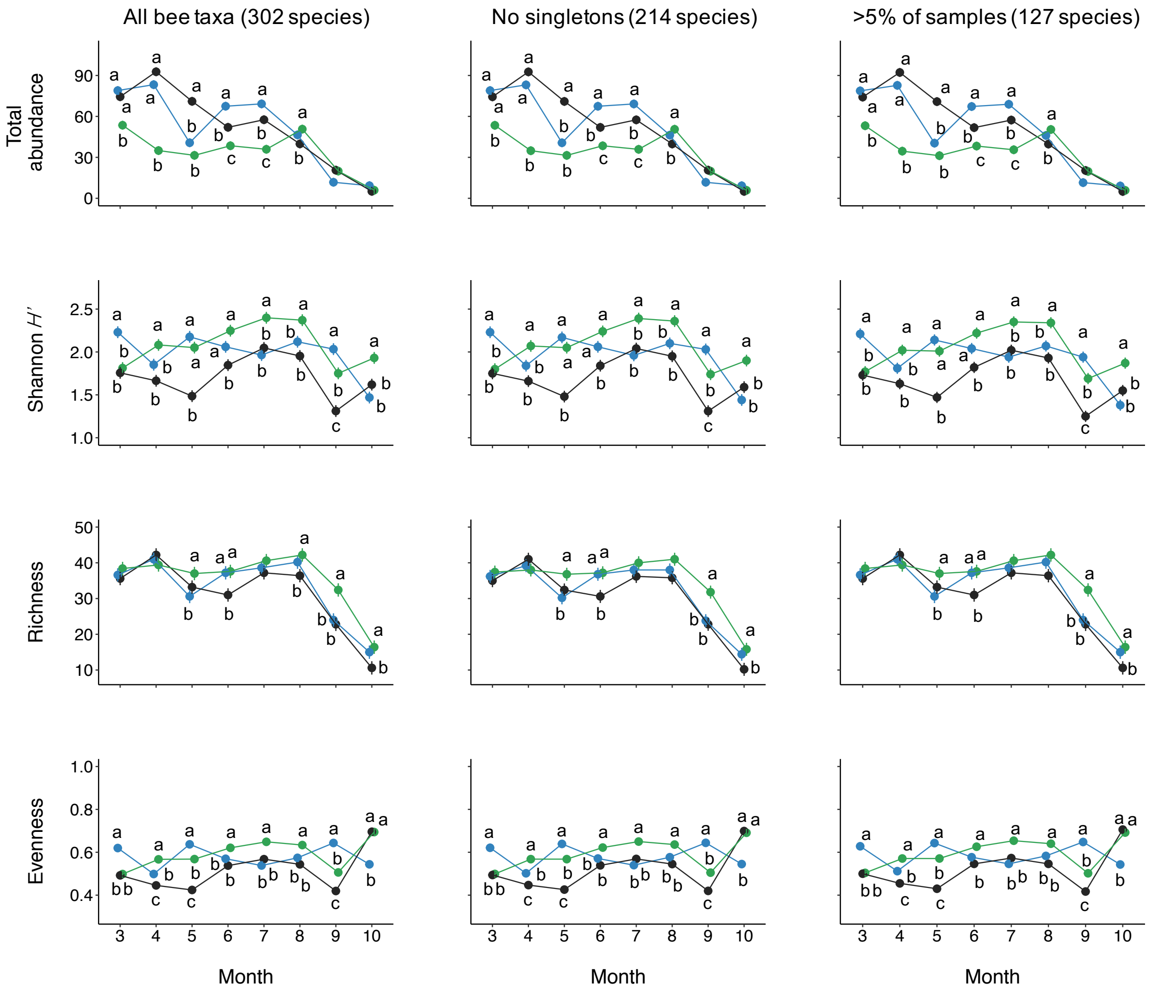


**Supplementary Figure S6.** Variation across sampling months in per-transect bee abundance, Shannon diversity (*H'*), richness, and evenness (Pielou's *J*), ± s.e., for three dryland ecosystems: Plains grassland (blue points), Chihuahuan Desert grassland (black points), and Chihuahuan Desert shrubland (green points). Analyses were run for the full dataset (left column), the dataset with singleton bee species removed (center column), and the dataset with only bee species present in >5% of samples (right column). Letters denote contrasts between ecosystems within a given month; ecosystems labeled with different letters differed significantly from one another in the relevant abundance/diversity metric. Points lacking letters did not differ significantly from any other ecosystem in the given month.
